# Acquired resistance to sotorasib in *KRAS*^*G12C*^ mutant NSCLC is vulnerable to PI3K-mTOR pathway inhibition mediated by 4E-BP1, a regulator of cap-dependent translation

**DOI:** 10.1101/2025.03.18.643954

**Authors:** Ismail M Meraz, Shuhong Wu, Yi Xu, Lihui Gao, Meng Feng, Chenghui Ren, Renduo Song, Ran Zhang, Qi Wang, Yuanxin Xi, Sung Yun Jung, Jing Wang, Bingliang Fang, Mourad Majidi, Jack A Roth

## Abstract

Sotorasib and adagrasib have shown significant efficacy in *KRAS*^*G12C*^ mutant NSCLC; however, acquired resistance (AR) occurs within 6–12 months. While some resistance arises from new mutations, over half of the resistant cases lack identifiable genomic alterations. We hypothesize that resistance is driven by signaling network rewiring, creating new therapeutic vulnerabilities. To investigate AR mechanisms, multiple AR models, including cell lines, PDXs, CDXs, and PDXOs, were developed. H23AR and H358AR cells displayed >600-fold and 200-fold; and PDXO303AR and PDXO314AR organoids exhibited >300-fold and >100-fold resistance to sotorasib, respectively; however, no additional mutations in *KRAS* or other potential genetic alterations were identified. The AR cells and PDXOs also showed comparable resistance to adagrasib. Distinct protein signatures associated with *KRAS* reactivation, mTORC1 signaling upregulation, and PI3K/AKT/mTOR pathway activation were identified in TC303AR & TC314AR PDXs. PI3K protein levels were significantly elevated in AR PDXs, H23AR, and H358AR cells. Pharmacological inhibition of PI3K with copanlisib or genetic knockout via CRISPR-Cas9 restored sotorasib sensitivity, suppressed colony formation, and inhibited downstream effectors, including p-AKT, p-mTOR, p-S6, p70S6K, p-GSK3β, and p-PRAS40 in AR cells. Copanlisib also sensitized both acquired and primary resistant PDXOs and synergized with sotorasib in restoring drug sensitivity. p4E-BP1 was significantly upregulated in H23AR and H358AR cells, which is suppressed by copanlisib. The level of p4E-BP1 expression was correlated with Sotorasib sensitivity in PI3K knockout clones, where the most sensitive clone displayed reduced or no p4E-BP1 expression. CRISPR-Cas9-mediated knockout of 4E-BP1, either alone or in combination with PI3K knockout, dramatically restored sotorasib sensitivity to levels comparable to parental cells. Suppression of 4E-BP1 hyperphosphorylation required dual inhibition of mTORC1 and mTORC2, and treatment with AZD8055 or sapanisertib (mTORC1/2 dual inhibitors) significantly dephosphorylated 4E-BP1 and restored sotorasib sensitivity in resistant cells and PDXOs. In PDX, CDX, and xenograft models in vivo, the combination of sotorasib with either copanlisib or sapanisertib resulted in robust, synergistic, and durable tumor regression at well-tolerated doses. These findings showed the critical role of PI3K/mTOR signaling as a bypass mechanism of resistance to *KRAS*^*G12C*^ inhibitors. We conclude that mTORC1/2 mediated inhibition of p4E-BP1 and combination strategies targeting this pathway effectively overcome acquired resistance to *KRAS*^*G12C*^ inhibitors in NSCLC.

## Introduction

*KRAS* mutations represent the most common gain-of-function alterations in cancer, accounting for approximately 30% of lung adenocarcinomas— a significantly higher prevalence than other oncogenic drivers, such as *EGFR* (15%), *ALK* rearrangements (5%), and *MET* alterations (3%), for which targeted therapies are available (1-3). Among *KRAS* mutations, the *KRAS*^*G12C*^ variant is one of the most frequent activating alterations, occurring in approximately 14% of lung adenocarcinomas and 0.5% to 4% of squamous cell carcinomas (4). This mutation is also found in 3% to 4% of colorectal cancers (CRC) and 1% to 2% of biliary and pancreatic cancers (5). Unlike protein kinases that utilize ATP as substrate, the KRAS protein is a GTPase that hydrolyzes guanosine triphosphate (GTP) to guanosine diphosphate (GDP) and normally functions as a molecular switch that cycles between an active, GTP–bound state and an inactive, GDP–bound form. Oncogenic *KRAS* mutations impair GTP hydrolysis, resulting in an accumulation of the active GTP-bound form, which drives pro-tumorigenic signaling predominantly through the mitogen-activated protein kinase (MAPK) and phosphatidylinositol 3-kinase (PI3K) pathways, promoting tumor growth and proliferation. Although inhibitors targeting several components of the MAPK and PI3K signaling pathways have been clinically approved, no direct *KRAS* inhibitors were available until recently. The attempts to use these drugs targeting downstream effectors in *KRAS* mutant cancers have largely been unsuccessful in the clinic (6). The recent discovery and development of new targeted therapeutic agents against *KRAS*^*G12C*^ renewed the clinical interest in *KRAS* biology, which was long considered undruggable for decades because of its high affinity for nucleotide and the lack of tractable binding pockets for small-molecule inhibitors (7,8).

Two FDA-approved small-molecule *KRAS*^*G12C*^ inhibitors, adagrasib (MRTX849) and sotorasib (AMG510), have shown potent, selective, and irreversible inhibition of *KRAS*^*G12C*^ activity, showing promising efficacy in non-small cell lung cancer (NSCLC) and more modest activity in colorectal cancer in early-phase clinical trials (9-11). In advanced NSCLC patients harboring *KRAS*^*G12C*^ mutations, both agents exhibited encouraging clinical outcomes, with objective response rates ranging between 37% and 45% and disease control rates of 81% to 96%, respectively (12). These inhibitors covalently bind to the switch II pocket of KRAS, which becomes accessible in the GDP-bound inactive state, irreversibly locking the protein in its inactive conformation (13). This mechanism inhibits downstream MAPK signaling, as evidenced by reduced ERK phosphorylation (p-ERK), and suppresses the viability of *KRAS*^*G12C*^-mutant cancer cell lines (9,10). Despite these promising results, approximately half of the patients treated with sotorasib or adagrasib in clinical trials did not achieve significant tumor shrinkage, highlighting the limited overall therapeutic benefit (14). Moreover, like other targeted therapies, virtually all patients who initially experience objective responses or stable disease eventually develop disease progression within 6–12 months of therapy initiation due to the emergence of resistance mechanisms. Therefore, the identification of these acquired resistance mechanisms, along with the development of alternative therapeutic strategies, including combination therapies with *KRAS* inhibitors, represents an urgent unmet clinical need.

Various potential mechanisms of acquired resistance to *KRAS*^*G12C*^ inhibitors have been identified in recent studies and can be broadly divided into the following categories: genetic alterations, reactivation of KRAS signaling, and non-genetic alternative bypass pathways (15). Genetic alterations, particularly the emergence of secondary KRAS mutations or new mutations in other oncogenic drivers, are often considered the primary cause of acquired resistance. However, these genetic events alone cannot fully explain treatment failure, as nearly half of patients treated with sotorasib or adagrasib did not show detectable mutations (16,17). Additionally, new secondary mutations in patients treated with KRAS inhibitors typically appear at low allelic frequencies, suggesting they may not be solely responsible for the observed tumor resistance (16-18). Reactivation of *KRAS* signaling has become a key mechanism of acquired resistance, mainly through the reactivation of two major canonical downstream pathways: the BRAF-MEK-ERK (MAPK) and PI3K-AKT-mTOR cascades. Besides these canonical pathways, several alternative bypass mechanisms have been identified that either lead to RAS pathway reactivation or activate parallel signaling pathways, such as the PI3K/mTOR axis, to sustain tumor growth and survival (16-20). Notably, inhibiting both MAPK and PI3K/mTOR pathways is often necessary to achieve significant antitumor effects, highlighting the importance of combination therapies to overcome resistance. Combination therapy is a critical strategy to prevent or delay resistance onset and to improve the effectiveness of *KRAS*^*G12C*^ inhibitors. Rational combination strategies that either target resistance mechanisms or synergize with *KRAS*^*G12C*^ inhibitors to suppress tumor growth and induce cancer cell death are urgently needed in clinical practice. A recent preclinical study demonstrated that combining a *KRAS*^*G12C*^ inhibitor with a selective mTORC1 kinase inhibitor produces a synergistic antitumor effect by inhibiting cap-dependent translation, a vital process for cancer cell survival and proliferation (21). Consistent with this, our recent research identified a non-genetic bypass resistance mechanism to osimertinib in *EGFR*-mutant NSCLC, where PDK1 (3-phosphoinositide-dependent kinase1) amplification drives resistance through upregulation of the AKT-mTOR signaling pathway (19).

Here, we take advantage of a series of paired sotorasib-resistant patient-derived xenografts (PDX), cell-derived xenografts (CDX), acquired resistant PDX-derived organoids (PDXO), and sotorasib-resistant isogeneic cell lines to identify acquired resistance mechanisms and explore potential therapeutic approaches. Reactivation of KRAS signaling was identified in acquired resistant tumors, which was evidenced by the impaired inhibition of ERK1/2 signaling. The upregulation of PI3K signaling in resistant cells was found as a major bypass mechanism for the acquired resistance. Genetic ablation of PI3K through CRISPR-mediated knockout or pharmacological inhibition with copanlisib restored sensitivity to both sotorasib and adagrasib in vitro and in vivo. The combination of PI3K inhibition with *KRAS*^*G12C*^ inhibitors exerted a synergistic antitumor effect by suppressing 4E-BP1 phosphorylation, thereby limiting cap-dependent protein translation, which was dependent on the simultaneous inhibition of both mTORC1 and mTORC2.

## Results

### Development and Characterization of Acquired Sotorasib-Resistant NSCLC PDX Models Harboring *KRAS* ^*G12C*^ Mutations

Among more than 200 NSCLC PDXs in our UT MD Anderson lung cancer PDX core, eleven well-characterized models harboring the *KRAS*^*G12C*^ mutation were identified. Whole-exome sequencing (WES) confirmed their co-mutation profiles in Table in **Figure 1A**. Four PDX models—TC247 (*KRAS*^*mt*^*/STK11*^*mt*^*/KEAP1*^*mt*^), TC303 (*KRAS*^*mt*^*/TP53*^*mt*^*/KEAP1*^*mt*^), TC314 (*KRAS*^*mt*^*/TP53*^*mt*^), and TC453 (*KRAS*^*mt*^*/STK11*^*mt*^)—were selected for evaluating sotorasib sensitivity based on their distinct co-mutation patterns. The selected models have co-mutations that are common in clinical specimens and are associated with drug resistance (15).

**Figure 1:**
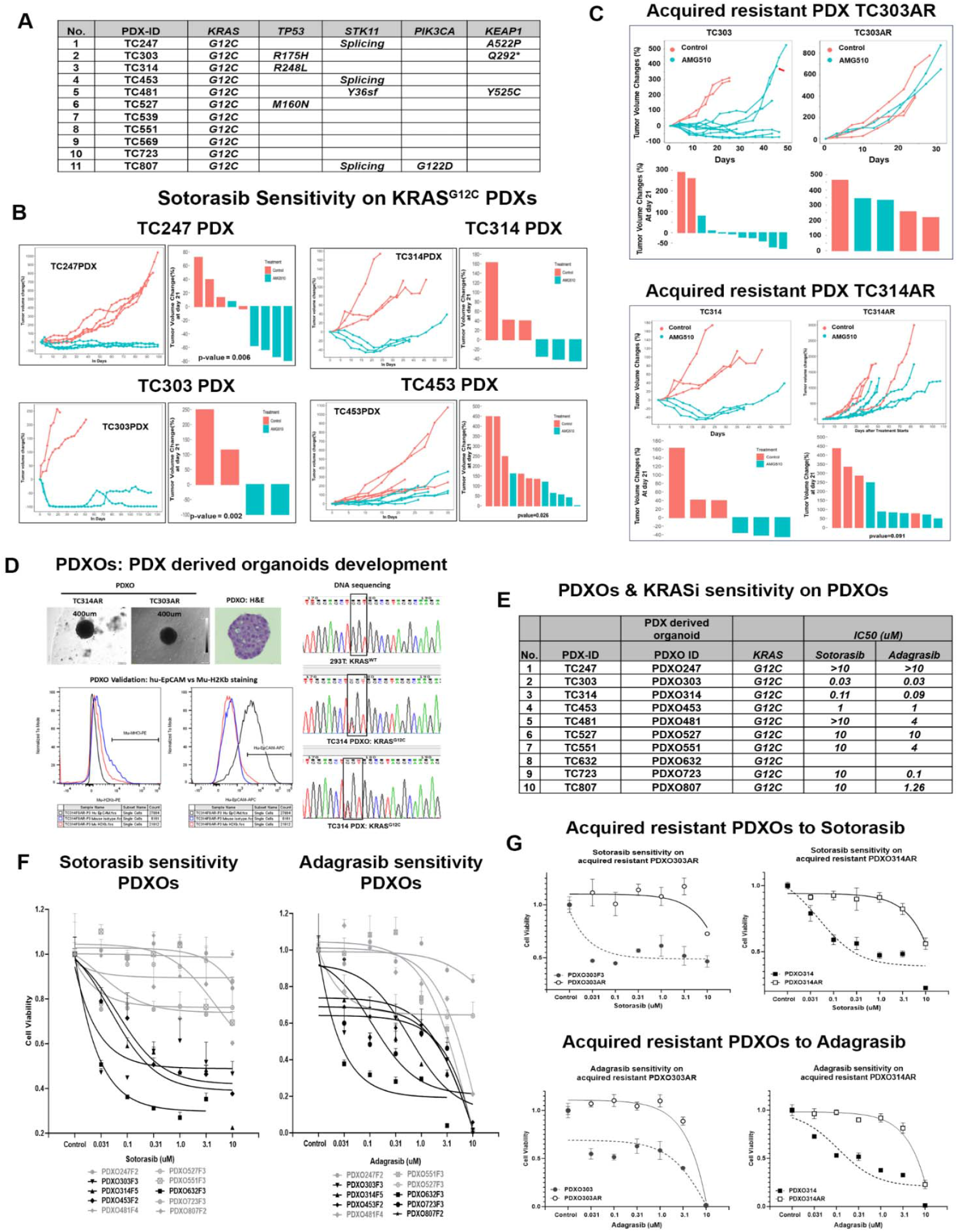
Development and characterization of sotorasib-acquired resistant patient-derived xenografts (PDXs) and PDX-derived organoids (PDXOs). **A)** A list of *KRAS*^*G12C*^ mutant-bearing NSCLC PDXs was developed in our PDX core. All the PDXs underwent whole-exome sequencing to identify their co-mutation status. **B)** Effect of sotorasib treatment on four *KRAS*^*G12C*^ mutant NSCLC PDXs (TC247, TC303, TC314, and TC453). PDX tumors were treated with sotorasib (100 mg/kg) when tumor volume reached around 200 mm^3^. Control mice were treated with the solvent. Sotorasib or solvent was administered QD orally for 21 days. Tumor volume changes in individual mice were calculated for treatment starting on day 0. **C)** Third-generation sotorasib-acquired resistant TC303AR and TC314AR PDXs were developed through continuous treatment of sotorasib for a long period of time, followed by passaging and reestablishing the PDX into new mice. The cycle of treatment and passaging was repeated three times. The effect of sotorasib treatment was evaluated on the 3rd-generation acquired resistant PDXs and compared with their parental TC303 and TC314 PDXs. **D)** PDXOs were developed and characterized. Whole exome sequence (WES) in PDXOs showed the KRAS^G12C^ mutation status (right panel), and the purity was determined by flow cytometry for detecting any mouse cell contamination by staining with mouse MHC Class I antibody (H2kb) and human epithelial marker (EpCAM) (left panel). **E)** The list of PDXOs developed from their respective PDXs and their sensitivity to sotorasib and adagrasib were determined based on IC50 (µM). **F)** Cell viability assay was performed on PDXOs for sotorasib (Left panel) and adagrasib (Right panel) sensitivity. Bold black lines indicate the PDXOs that were sensitive, and ash color lines indicate the PDXOs that were resistant to KRAS inhibitors. **G)** Sotorasib-acquired resistant PDXO303AR and PDXO314AR were developed through repeated cycles of sotorasib treatment and passaging of PDXOs until they developed resistance, and these acquired resistant PDXOs were then treated with sotorasib (top panel) and adagrasib (bottom panel), and the sensitivity was compared with their parental counterparts (PDXO303 & PDXO314). Data shown represent the mean□±□SE of three independent experiments.

To investigate the sotorasib sensitivity in vivo, these PDXs were engrafted in NSG mice and treated with sotorasib when tumors reached ∼200 mm^3^. Despite co-mutations in *STK11, TP53, or KEAP1*, all four models exhibited high sensitivity to sotorasib, leading to significant tumor growth suppression (**Figure 1B**). Notably, TC247, TC303, and TC314 showed substantial tumor regression, whereas TC453 displayed only moderate sensitivity without significant regression.

To generate sotorasib-resistant models, the two most sensitive PDXs (TC303 and TC314) underwent prolonged in vivo treatment. Third-generation (G3) resistant PDXs—TC303AR and TC314AR—were established by re-passaging the PDXs into new mice, followed by sotorasib treatment for at least 3 cycles. These resistant models no longer responded to sotorasib (**Figure 1C**).

WES of TC303AR and TC314AR confirmed that both retained the *KRAS*^*G12C*^ mutation, with no additional mutations detected in *KRAS (***Figure 1-figure supplement 1**).

### Development and Characterization of Acquired Resistance to Sotorasib in NSCLC PDX-Derived Organoids

Ten PDX-derived organoids (PDXOs) listed in the table in **Figure 1E** were established and characterized from their corresponding PDX models. All organoids were characterized by sequencing and flow cytometry. WES confirmed that all established organoids retained the *KRAS*^*G12C*^ mutation, while flow cytometry analysis detected no mouse cell contamination, as indicated by the absence of H2kb staining (mouse MHC class I) and the presence of human EpCAM, an epithelial marker (**Figure 1D**).

Sotorasib sensitivity was assessed for all PDXOs, with IC50 values (µM) listed in the table shown in **Figure 1E**. The sensitivity testing revealed that PDXO303 and PDXO314 were the most responsive, with IC50 values of 0.03 µM and 0.11 µM, respectively. These two organoids were also challenged for Adagrasib sensitivity. Both models also exhibited high sensitivity to adagrasib, the second FDA-approved *KRAS*^*G12C*^ inhibitor.

Of the remaining PDXOs, six out of seven displayed primary resistance to sotorasib (IC50 ≥10 µM), with only one of the seven showing sensitivity to adagrasib. Specifically, PDXO303, PDXO314, and PDXO723 were highly sensitive to adagrasib (IC50 0.03 µM, 0.09 µM, and 0.1 µM, respectively, **Figure 1E & F**).

PDXO303 and PDXO314, the two most sensitive PDXOs to both sotorasib and adagrasib, underwent prolonged exposure to increasing doses of sotorasib in culture. This resulted in the development of sotorasib-resistant PDXO303AR and PDXO314AR, exhibiting >300-fold and >100-fold resistance, respectively. Their IC50 values increased from 0.03 µM to >10 µM (PDXO303AR) and 0.11 µM to >10 µM (PDXO314AR) (**Figure 1G**). Both resistant models also displayed significant resistance to adagrasib.

To verify the KRAS mutation status, WES was performed for the resistant PDXOs. Both PDXO303AR and PDXO314AR retained the *KRAS*^*G12C*^ mutation, and no additional mutations were detected in *KRAS*.

### Development of Acquired Sotorasib-Resistant NSCLC Isogenic Cell Lines and Their Sensitivity to *KRAS*^*G12C*^ Inhibitors

Two NSCLC cell lines (H23 and H358) harboring the *KRAS*^*G12C*^ mutation were selected to develop acquired resistance. Cells were continuously exposed to increasing doses of sotorasib until they showed sustained growth in the presence of 2.5 µM (H23) and 1 µM (H358). The resulting resistant cell lines, H23AR and H358AR, exhibited over 600-fold and 200-fold resistance, respectively (**Figure 2A**).

**Figure 2:**
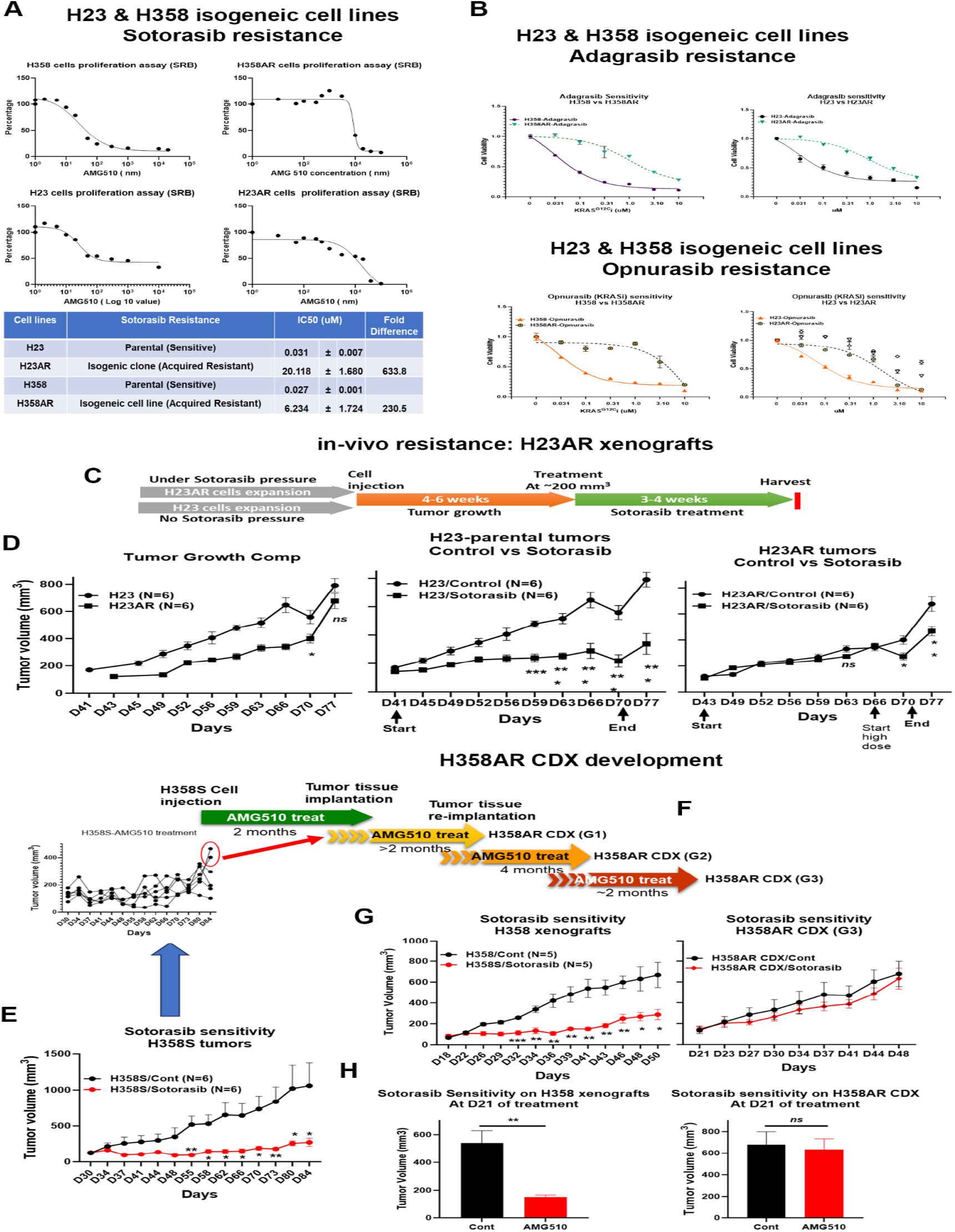
Evaluation of sotorasib resistance on sotorasib-acquired resistant isogeneic cell lines and acquired-resistant cell line-derived xenograft (CDX). **A)** SRB (Sulforhodamine B) assay was performed to determine the cell viability after sotorasib treatment, and the sotorasib sensitivity was compared between H23 (parental) vs H23AR (resistant) cells and H358 (parental) vs H358AR (resistant) cells. IC50 values for sotorasib were determined, and the fold of resistance is listed in the table (bottom panel). **B)** Cell viability assay showed the sensitivity of H23 & H23AR and H358 & H358AR isogeneic cells to adagrasib (top panel) and Opnurasib (bottom panel), two other KRAS^G12C^-specific inhibitors. **C)** The strategy for the evaluation of the in vivo resistance to sotorasib for H23AR xenograft tumors is shown. **D)** H23 and H23AR cells were implanted into mice and treated with sotorasib (100mg/kg) for at least 3 weeks when tumor sizes reached approximately 200 mm^3^. H23AR cells were expanded in culture under sotorasib pressure before implantation. The antitumor effect of sotorasib was evaluated in both H23 xenograft (control tumor vs treated tumors) and H23AR xenograft tumor models (control vs sotorasib). Statistical significance was calculated. **E)** Development of acquired resistant H358AR CDX. First, H358 xenograft tumors were developed from parental H358 cells and treated with sotorasib at a daily dose of 100 mg/kg for an extended period, as shown in the figure. The antitumor effect of sotorasib on H358 parental tumors was evaluated, and their statistical significance was calculated. **F)** Second, the residual tumors that developed initial resistance were passaged into new mice and treated again with another cycle of sotorasib treatment when tumors reached around 200mm^3^. The cycle of passaging and treatment was repeated three times at the indicated times shown in the figure until the third generation of acquired resistant H358AR CDXs was developed. The strategy shown here for the development of sotorasib-acquired resistant H358AR CDX. **G)** Third-generation sotorasib-resistant H358AR CDX showed no significant effect with sotorasib treatment, whereas H358 parental xenograft tumors showed significant sensitivity to sotorasib. **H)** Sotorasib antitumor effect was calculated based on tumor volume shrinkage after 21 days of treatment, where H358AR CDX showed no significant effect of sotorasib treatment. Each experiment was repeated three times. Statistics are shown at a significance level of p<0.05 unless otherwise noted. Data is shown as mean percentage ±SD, n>5 mice/group was used in each experiment. *, *P* < 0.05; **, *P* <0.005; *** *P* <0.0005.

Both resistant lines were also tested for sensitivity to adagrasib and opnurasib, two additional *KRAS*^*G12C*^ inhibitors. Like sotorasib, H23AR, and H358AR displayed high resistance to both inhibitors, with no significant cell death observed (**Figure 2B**).

Whole-exome sequencing (WES) confirmed the retention of the *KRAS*^*G12C*^ mutation in both resistant cell lines, with no additional mutations detected in *KRAS*. However, H23AR acquired 23 SNPs and 3 indels, while H358AR acquired 82 SNPs and 7 indels compared to their sotorasib-sensitive counterparts (**Figure 2-figure supplement 1**).

### In Vivo Evaluation of Sotorasib Resistance in NSCLC Xenograft Models Derived from Acquired-Resistant H23AR Cells

To evaluate the in vivo sensitivity of sotorasib in acquired resistant cells, H23AR cells, which exhibited over 600-fold resistance in vitro, were implanted into mice to generate xenograft tumors. Before implantation, H23AR cells were expanded in culture under sotorasib pressure to maintain resistance.

Both H23 (parental) and H23AR xenografts were established, and sotorasib treatment was initiated when tumors reached approximately 200 mm^3^ (treatment strategy shown in **Figure 2C**). Although there were significant differences in tumor growth between H23 and H23AR tumors, however, as expected, sotorasib exerted a significant antitumor effect on H23 parental tumors, leading to tumor growth suppression. In contrast, tumors derived from H23AR cells showed no tumor growth inhibition, confirming that H23AR cells retained their resistance in vivo (**Figure 2D**).

### Development of Sotorasib Acquired-Resistant H358 Cell Line-Derived Xenograft Model

The H358 cell line, which harbors the *KRAS*^*G12C*^ mutation, is among the most sotorasib-sensitive NSCLC cell lines, with an IC50 of 27 nM. To develop a sotorasib-acquired resistant cell line-derived xenograft (CDX) model, xenograft tumors were established using parental H358 cells and treated daily with sotorasib for two months. Tumors in the sotorasib-treated group exhibited a significant reduction in growth compared to the vehicle-treated controls (**Figure 2E**).

Following two months of treatment, residual tumor tissues were re-implanted into new mice and subjected to another two-month sotorasib treatment cycle. This process was repeated for three cycles, ultimately generating a third generation (G3) sotorasib-resistant H358AR CDX (**Figure 2F**). By the third generation, H358AR CDX tumors no longer exhibited significant antitumor responses to sotorasib (**Figure 2G**).

To confirm the extent of sotorasib resistance, parental H358 xenografts and H358AR CDX (G3) tumors were implanted and subsequently treated with sotorasib. After 21 days of treatment, parental H358 tumors remained highly sensitive, whereas H358AR CDX tumors displayed minimal to no response (**Figure 2H**).

Whole-exome sequencing (WES) confirmed that H358AR CDX tumors retained the *KRAS*^*G12C*^ mutation, with no additional KRAS mutations detected.

### Reactivation of KRAS and upregulation of PI3K signaling in acquired sotorasib-resistant PDX models

To investigate the mechanisms underlying sotorasib-acquired resistance, an unbiased mass spectrometry (MS)--based proteomics approach was utilized to profile the global and phosphoproteome of sotorasib-resistant TC303AR and TC314AR isogenic PDXs. The global and phosphoproteomic analyses identified over 8,000 and 4,000 gene protein products (GPPs), respectively. Samples were analyzed using label-free nanoscale liquid chromatography coupled with tandem mass spectrometry (nanoLC-MS/MS) on a Thermo Fusion Mass Spectrometer. The resulting data were processed and quantified using the Proteome Discoverer 2.5 interface with the Mascot search engine, referencing the NCBI RefSeq protein database (Saltzman, Ruprecht).

Dendrogram clustering showed that TC303 and TC314 PDX samples formed distinct clusters, highlighting their biological differences. Within each model, parental and resistant PDXs clustered separately, confirming proteomic alterations associated with sotorasib resistance (**Figure 3A**). A two-component analysis further distinguished sensitive from resistant PDXs, while also differentiating TC303 from TC314 PDX models (**Figure 3B**).

**Figure 3:**
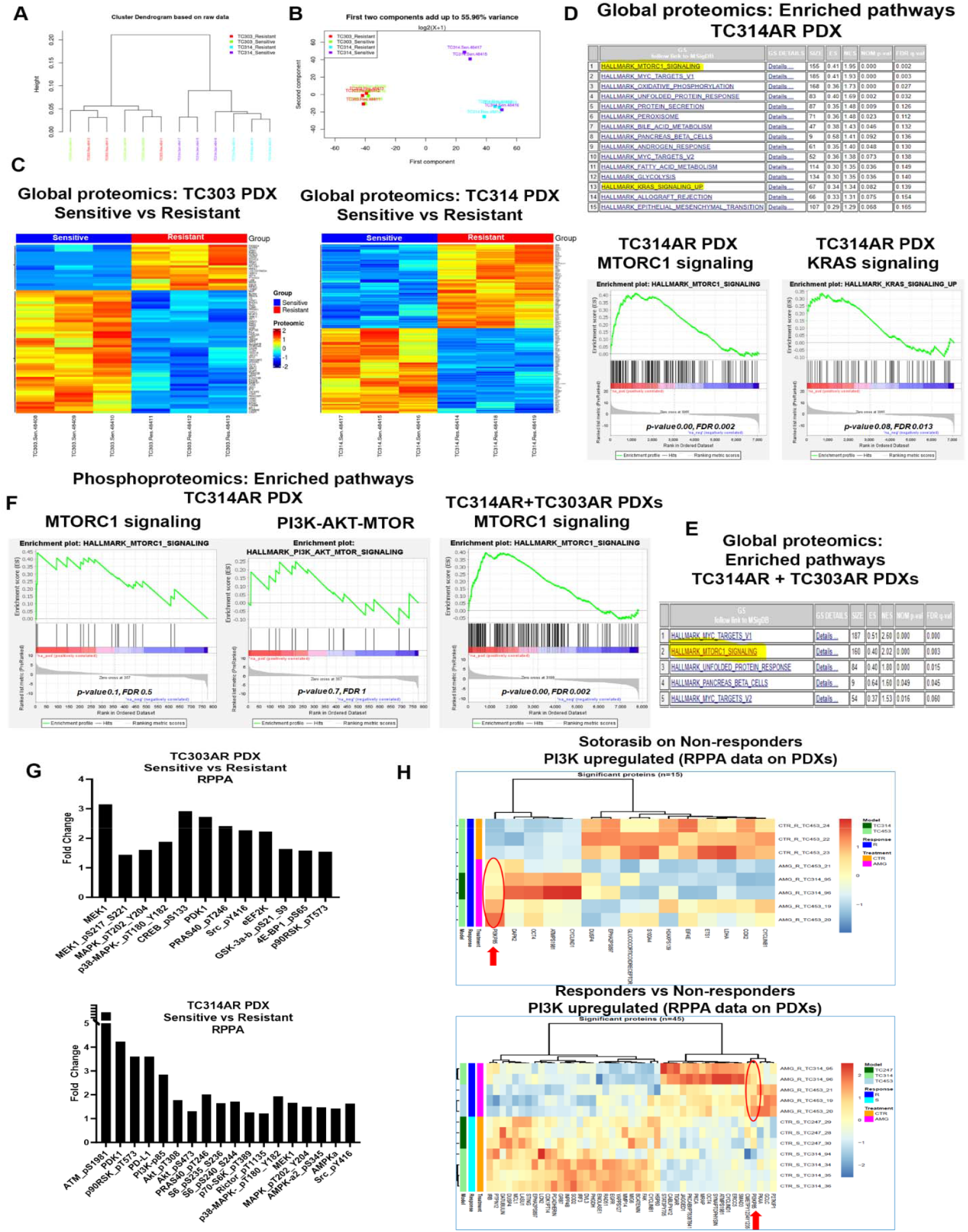
Reactivation of KRAS and Upregulation of PI3K/AKT/mTOR Signaling in Acquired Sotorasib-Resistant PDX Models. Acquired resistant TC303AR and TC314AR PDXs and their isogeneic TC303 and TC314 PDXs snap frozen samples underwent global and phospho-proteomics by mass spectrometry. The data were analyzed in the Biostatistics Department at the MD Anderson Cancer Center. **A-B)** Dendrogram plot and two-component analysis curve showed the clustering of the samples, indicating the similarities and distances among samples. **C)** Heat-map was generated among the proteins that were statistically significantly upregulated or downregulated between parental TC314 and resistant TC314AR PDX samples (right panel) and parental TC303 vs resistant TC303AR PDX samples (left panel). **D)** Pathway Enrichment analysis was performed among significantly altered global proteins in TC314AR samples. A list of gene-sets enriched was shown in the top panel with their FDR values, and the enrichment of gene-sets associated with mTORC1 and KRAS signaling is shown in the bottom panel with their significant p-value and FDR. **E)** Enrichment analysis was performed for global proteomics for combined TC303AR and TC314AR data. A list of enriched gene-sets was listed in the right panel, and the mTORC1 signaling enrichment graph was shown in the left panel. **F)** Pathway Enrichment analysis in phosphoproteomics in TC314AR PDX was performed. The enriched mTORC1 signaling is shown in the left panel, and the PI3K/AKT/mTOR pathway enrichment is shown in the right panel. FDR and p-values are displayed in the graphs. **G)** RPPA data derived from TC303AR and TC314AR isogenic PDX samples, and the significantly altered proteins associated with MAPK and PI3K/AKT/mTOR pathways were analyzed and compared. The increased expression by fold-change of these proteins between TC303 parental vs TC303AR resistant samples (top) and TC314 parental vs TC314AR resistant samples (bottom) was shown. **H)** PDXs TC247, TC314, & TC453 were implanted into mice, and when the PDX tumor volume reached 200 mm^3^, tumors were treated with sotorasib (100 mg/kg) for 3 weeks. After 21 days of treatment, the treatment efficacy was determined based on the percentage of tumor growth inhibition (details in the method section) and categorized between responders and non-responders. PDX samples underwent RPPA analysis. Heat-maps (bottom panel) were generated based on the significantly altered proteins and compared between responder vs non-responder groups. Non-responder PDXs were grown again and treated with sotorasib for 21 days. After treatment, tumors were harvested, and RPPA was performed. The RPPA data derived from sotorasib-treated non-responders and control non-responders were compared, and a heat-map was generated among significantly altered proteins between control vs treated among non-responder PDXs (top panel). N≥3 biological replicates were used for each PDX for Mass Spectrometry. N=5 PDX-tumors/treatment group, and N ≥ 3 PDX samples from each treatment group were used for RPPA analysis. The criteria of protein selection for significant up- or down-regulation were: 1. Significant in overall F-test (FDR-adjusted p-value<0.05); 2. Significant in pairwise comparison. (FDR-adjusted p-value<0.05).

Global proteomic heatmap analysis revealed unique protein expression profiles in TC303AR and TC314AR PDXs compared to their sensitive counterparts (**Figure 3C**). A distinct set of proteins was upregulated, and another set of proteins was downregulated in TC303AR PDX which were completely different than their sensitive counterpart (**Figure 3C; left**). Similarly, a distinct set of global proteins was significantly altered in TC314AR PDX as compared with sensitive TC314 PDX (**Figure 3C; right**) (Heatmap cutoff p-value <0.01). A total of 293 proteins in TC303AR and 535 proteins in TC314AR were significantly altered as compared with their respective sensitive counterpart. A Venn diagram showed that only 32 proteins (∼4%) were commonly dysregulated in both resistant models. Notably, 65% of altered proteins were unique to TC314AR, whereas 33% were distinct to TC303AR (**Figure 3-figure supplement 1**).

Pathway enrichment analysis of TC314AR global proteomics identified 26 upregulated gene sets, of which 15 were significant at FDR <25%, 10 at nominal p <5%, and 5 at nominal p <1%. MTORC1 signaling emerged as one of the most upregulated pathways in TC314AR PDXs (FDR q <0.001), along with *KRAS* signaling (nominal p <0.08) when compared with their sensitive counterpart, TC314 (**Figure 3D**). When enrichment analysis was performed across both resistant PDX models (TC303AR + TC314AR vs TC303 + TC314), MTORC1 signaling remained one of the most upregulated pathways (FDR q <0.003) (**Figure 3E**).

Further phosphoproteomic analysis in TC314AR PDXs revealed enhanced activation of the MTORC1 and PI3K-AKT-mTOR pathways as compared with the TC314-sensitive PDX, suggesting a potential adaptive resistance mechanism (**Figure 3F**). Upregulation of mTORC1 and PI3K/AKT/mTOR signaling was found in the phosphoproteomic analysis in TC303AR data, as well as in TC314AR+TC303AR combined data (**Figure 3-figure supplement 2**).

### Upregulation of PI3K-AKT-mTOR Signaling in Sotorasib-Resistant PDX Models

Reverse Phase Protein Array (RPPA) analysis in sotorasib-resistant TC314AR and TC303AR PDX models showed a significant upregulation of proteins associated with the PI3K-AKT-mTOR signaling pathway compared to their sensitive counterparts. Specifically, the expression levels of PI3K-p85, p-Akt, PDK1, p-GSK-3β, p-S6, p-PRAS40, p-4E-BP1 in the PI3K-AKT-mTOR pathway, as well as the expression level of MEK1, p-MEK1, p-MAPK, p-p38-MAPK, p-AMPK, and p-p90RSK in the MAPK pathway were markedly increased in resistant TC303AR and TC314AR PDXs (**Figure 3G**).

To further investigate the role of PI3K signaling in sotorasib response, RPPA analysis was performed on residual tumors from sotorasib-responding and non-responding PDX models. *KRAS*^*G12C*^-mutant PDXs (TC247, TC314, TC453) were implanted into mice to generate PDX tumors and treated with sotorasib for three weeks. Heatmap analysis revealed that PI3K expression was significantly elevated in non-responding PDX tumors, whereas it was downregulated in sotorasib-responsive tumors (**Figure 3H, lower panel**).

Furthermore, PI3K expression was found to be upregulated in sotorasib-treated vs. control tumors among non-responding PDXs, suggesting that PI3K signaling activation is associated with resistance to sotorasib treatment (**Figure 3H, upper panel**). These findings indicate that PI3K-AKT-mTOR pathway activation may play a critical role in mediating sotorasib resistance in *KRAS*^*G12C*^-mutant NSCLC and could represent a potential therapeutic target for overcoming resistance.

### KRAS Reactivation in Sotorasib-Resistant Cells via ERK Phosphorylation

KRAS inhibition is commonly assessed by evaluating the inhibition of ERK phosphorylation (p-ERK) (9). To confirm KRAS reactivation in acquired resistance, p-ERK levels were analyzed following sotorasib treatment. In parental H23 and H358 cells, which are highly sensitive to sotorasib, treatment resulted in significant inhibition of p-ERK, indicating effective KRAS suppression (**Figure 3-figure supplement 3A-B**). However, in acquired resistant H23AR and H358AR cells, p-ERK levels remained unaffected by sotorasib treatment, suggesting KRAS reactivation and continued downstream signaling despite drug exposure (**Figure 3-figure supplement 3A-B**).

### Pharmacological inhibition of PI3K signaling by copanlisib sensitizes sotorasib-resistant cells and PDXOs

To assess the role of PI3K in acquired resistance, copanlisib, a pan-PI3K inhibitor, was tested on resistant cells and PDX-derived organoids (PDXOs). Treatment with copanlisib showed that both H23 and H23AR cells were highly sensitive, with IC50 values of 0.256 ± 0.038 µM and 0.394 ± 0.029 µM, respectively (**Figure 4A**). Similarly, H358 and H358AR cells showed high sensitivity to copanlisib, with IC50 values of 0.083 ± 0.011 µM and 0.063 ± 0.009 µM, respectively (**Figure 4A**).

**Figure 4:**
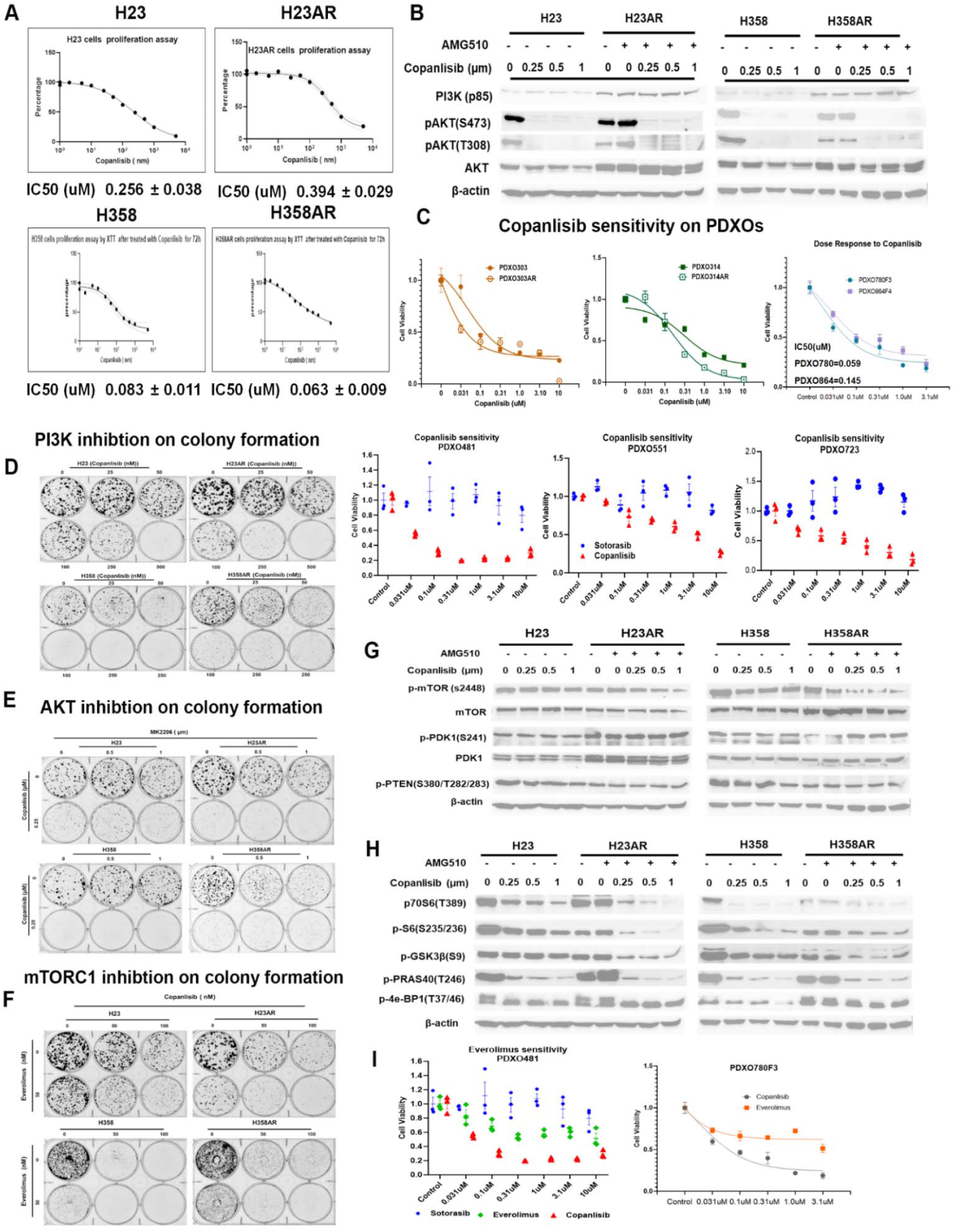
Pharmacological inhibition of PI3K signaling by copanlisib inhibits colony formation and downregulates PI3K/AKT/mTOR signaling molecules in acquired resistant cells and PDXOs. **A)** SRB (Sulforhodamine B) assay was performed to compare the copanlisib sensitivity between H23 (parental) vs H23AR (resistant) cells and H358 (parental) vs H358AR (resistant) cells. Both acquired resistant H23AR and H358AR were cultured and maintained under sotorasib pressure. For the experiment, cells were plated for copanlisib treatment at different doses for 72h, and the cell viability assay was performed by using the SRB method. The IC50 values of copanlisib for each cell line were determined and shown at the bottom of each respective graph. **B)** H23AR and H358AR cells were cultured in sotorasib-containing media and plated for copanlisib treatment at different doses for 24h. For no treatment control samples, H23AR and H358AR were omitted from sotorasib for the last 24h; otherwise, acquired cells were always maintained under sotorasib pressure. Western blot was performed for PI3K (p85), AKT, and phospho-AKT (S473 & T308). **C)** Effect of copanlisib on acquired resistant PDXOs. Acquired resistant PDXO303AR and PDXO314AR isogeneic organoids and primary resistant organoids (PDXO780 & PDXO664) were grown in 96-well plates according to the organoid protocol and treated with different doses of copanlisib. After the treatment, cell viability was assessed by the Cell-Glow assay. Cell viability of copanlisib was also compared with sotorasib on the sotorasib primary resistant organoids PDXO481, PDXO551, and PDXO723 (bottom panel). **D)** Inhibition of colony proliferation by copanlisib treatment at different doses on H23AR and H358AR isogeneic cells was shown in the colony formation assay. **E)** Effect of copanlisib and MK2206, and **F)** copanlisib and everolimus combination on the inhibition of cell proliferation in both resistant H23AR and H358AR cells, shown in colony formation assays. **G-H)** H23AR and H358AR cells were cultured, maintained, and seeded in media containing sotorasib, and treated with copanlisib doses mentioned in the figure for 24h. For no treatment control samples, H23AR and H358AR were omitted from sotorasib for the last 24h; otherwise, acquired cells were always maintained under sotorasib pressure. Western blot was performed for upstream **(G)** mTOR, p-mTOR, PDK, p-PDK, and p-PTEN; downstream **(H)** p70S6, p-S6, p-GSK-3b, p-PRAS40, p-4E-BP1 molecules. **I)** Cell viability assay on sotorasib primary resistant PDXO481 organoid was performed to compare the sensitivity among sotorasib, copanlisib, PI3K inhibitor, and everolimus, mTORC1 inhibitor (Left), and the sensitivity between copanlisib and everolimus was compared in sotorasib primary resistant PDXO780 organoid (Right). Each experiment was repeated three times. Data is shown as mean percentage ±SD, n=3.

Upregulation of PI3K signaling was observed in sotorasib-resistant PDXs, with a significant increase in PI3K (p85) expression in H23AR and H358AR cells compared to their respective parental counterparts (**Figure 4B**).

Copanlisib efficacy was also tested in sotorasib-resistant organoids, PDXO303AR and PDXO314AR, where copanlisib was effective in sotorasib-resistant organoids (**Figure 4C**). Notably, both parental and resistant PDXO303 and PDXO314 organoids exhibited comparable sensitivity to copanlisib, indicating its strong therapeutic potential.

Further, copanlisib sensitivity was evaluated in additional *KRAS*^*G12C*^-mutant PDXOs that were intrinsically resistant to sotorasib. PDXO780 and PDXO864 organoids were highly sensitive to copanlisib, with IC50 values of 0.059 µM and 0.145 µM, respectively (**Figure 4C**). Moreover, PDXO481, PDXO551, and PDXO723 organoids, which did not respond to sotorasib treatment, exhibited remarkable sensitivity to copanlisib (**Figure 4C**).

Expression of PI3K was upregulated in both H23AR and H358AR cells compared to their sensitive counterparts. Treatment with copanlisib effectively inhibited PI3K signaling, as evidenced by a significant reduction in AKT phosphorylation at S473 (p-AKT S473) and T308 (p-AKT T308) in both H23AR and H358AR cells (**Figure 4B**). Similarly, copanlisib treatment led to a marked decrease in AKT phosphorylation in parental H23 and H358 cells.

To evaluate the functional impact of PI3K inhibition, a colony formation assay was performed. H23 and H358 isogenic cells were treated with increasing concentrations of copanlisib (25 nM, 50 nM, 100 nM, 250 nM, and 500 nM). Even at 50 nM, copanlisib reduced colony formation in both resistant cell lines, with a significant inhibitory effect observed at 100 nM (**Figure 4D**).

To further investigate the role of AKT signaling, resistant cells were treated with MK2206, an AKT inhibitor, alone and in combination with copanlisib. MK2206 alone (0.5 µM) moderately reduced colony formation and its combination with copanlisib (250 nM) resulted in a significantly enhanced inhibitory effect, indicating a strong combinatorial effect on sensitizing sotorasib-resistant cells (**Figure 4E**).

Previous analyses showed significant enrichment of mTORC1 signaling in sotorasib-resistant PDXs (**Figure 3**). To investigate the role of mTORC1 inhibition in resistant cells, H23, H23AR, H358, and H358AR cells were treated with everolimus, an mTORC1 inhibitor. Everolimus inhibited colony formation in both resistant cell lines at a concentration of 50 nM. Notably, the combination of everolimus with a low dose of copanlisib (50 nM) resulted in a profound reduction in colony formation (**Figure 4F**). These findings suggest that PI3K inhibition with copanlisib is effective for sotorasib-resistant cells, although the drug is equally effective on the parental cells, highlighting the critical role of PI3K signaling in acquired resistance. The specific role of PI3K is shown in the knockout experiment in Figure 5.

**Figure 5:**
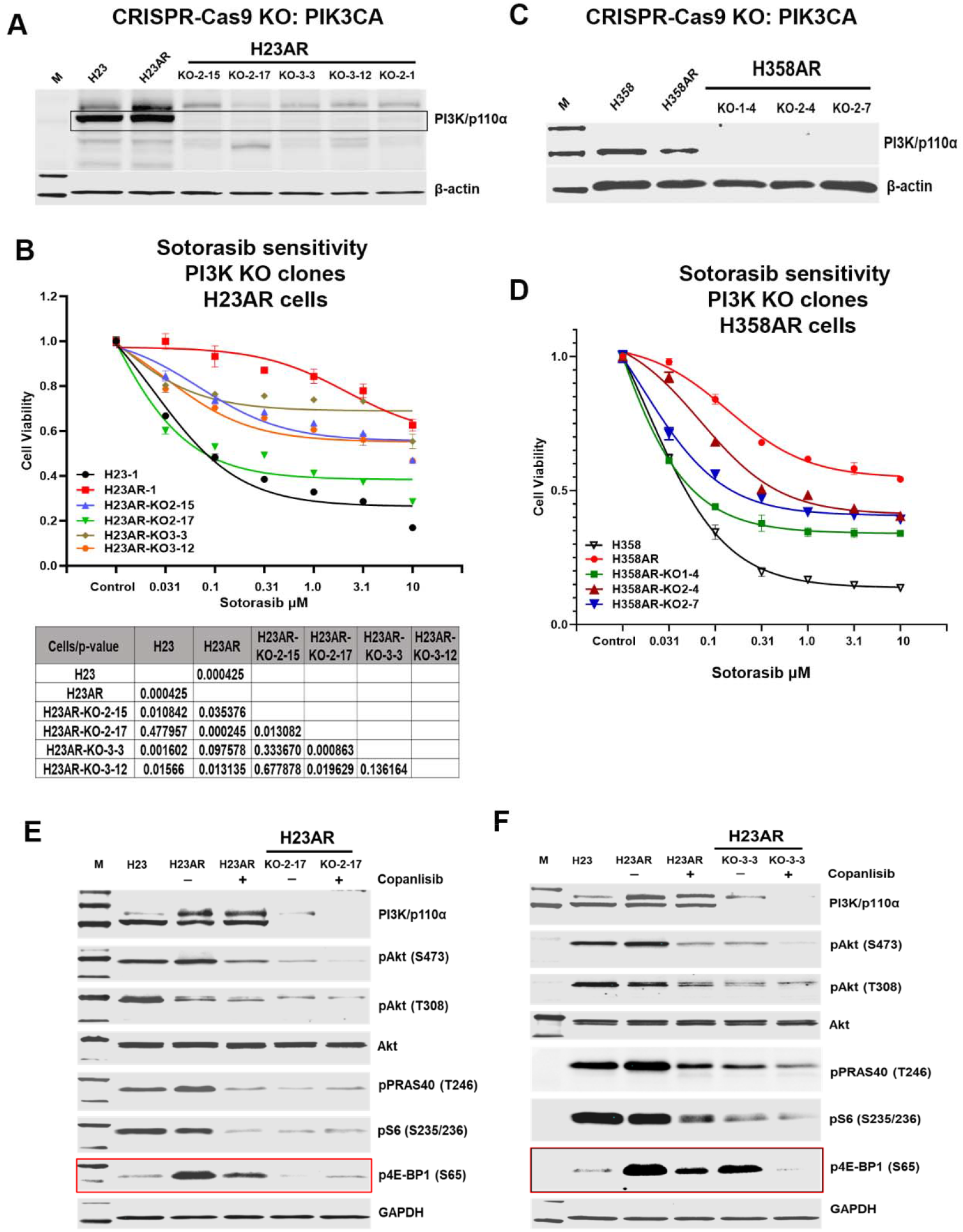
PI3K knockout via CRISPR-Cas9 restores sotorasib sensitivity in acquired resistant cells by dephosphorylating 4E-BP1. **A)** The PI3K/p110α subunit was knocked out using CRISPR-Cas9 technology on H23AR. Five clones, H23AR^*PI3K-KO-2-15*^, H23AR^*PI3K-KO-2-17*^, H23AR^*PI3K-KO-3-3*^, H23AR^*PI3K-KO-3-12*^ & H23AR^*PI3K-KO-2-1*^ from H23AR cells were selected, and the PI3K expression on these clones was verified by western blots. **B)** Cell viability assay was performed on these PI3K knockout clones generated from H23AR cells to compare the sotorasib sensitivity to determine the most sensitive and least sensitive clone. The sotorasib sensitivity on PI3K clones was also compared with the parental H23 and resistant H23AR cells. The statistical significance was compared among the clones and H23AR isogeneic cells, as shown in the table below the graph. **C)** The PI3K/p110α subunit was knocked out by CRISPR-Cas9 technology on H358AR. Three clones, H358AR^*PI3K-KO-1-4*^, H358AR^*PI3K-KO-2-4*^, and H358AR^*PI3K-KO-2-7*^ from H358AR cells were selected, and the knockout status of PI3K expression on these clones was verified by western blots. **D)** Cell viability assay was performed on these PI3K knockout clones generated from H358AR cells to compare the sotorasib sensitivity to H358AR-resistant and H358 parental cells. **E)** For the molecular analysis, H23, H23AR, and the most sensitive PI3K knockout clone H23AR^*PI3K-KO-2-17*^ were seeded and treated with copanlisib for 24h, and the downstream PI3K, AKT, p-AKT, p-S6, and p-4E-BP1 status was determined by the western blot. The effect of copanlisib treatment on downstream molecules and the level of expression was also compared among PI3K knockout and H23AR isogeneic cells. F) Similarly, for the molecular analysis, H23, H23AR, and the least sensitive PI3K knockout clone H23AR^*PI3K-KO-3-3*^ were seeded and treated with copanlisib for 24h, and the downstream PI3K, AKT, p-AKT, p-S6, and p-4E-BP1 status was determined by the western blot. The effect of copanlisib treatment on downstream molecules and the level of expression was also compared among PI3K knockout and H23AR isogeneic cells, as shown. Each experiment was repeated three times. Data is shown as mean percentage ±SD, n ≥ 3.

### Copanlisib modulates PI3K-AKT-mTOR signaling in sotorasib-resistant cells by regulating downstream molecules

PI3K-AKT-mTOR signaling was found to be a major signaling pathway upregulated in resistant cells. Inhibition of PI3K by copanlisib sensitizes the resistant cells. Here, we investigated the effect of copanlisib on various upstream and downstream molecules in the PI3K-AKT-mTOR pathway associated with copanlisib sensitivity. H23 & H23AR and H358 & H358AR cells were treated with multiple doses of copanlisib, and western blots were performed to evaluate the expression of upstream and downstream molecules associated with PI3K signaling.

Phosphorylation of mTOR at serine 2448 (s2448) was significantly inhibited by copanlisib treatment in both resistant H23AR and H358AR cells as compared to their respective sensitive parental cells, although the total level of mTOR remained unchanged (**Figure 4G**). Quantitative analysis was performed to confirm these findings (**Figure 4-figure supplement 1**). The levels of total PDK and phospho-PDK (S241) were increased in H23AR cells compared with H23, and neither was suppressed by copanlisib treatment in the resistant lines (**Figure 4G**). This pattern was confirmed by quantitative analysis of the western blot band intensities (**Figure 4—figure supplement 2**). The basal level of pPTEN was decreased in both resistant cells (H23AR vs H23 and H358AR vs H358), and the level was not altered by copanlisib treatment (**Figure 4G**).

Alterations by copanlisib treatment on various downstream molecules in the PI3K-AKT-mTOR pathway were assessed in both resistant cell lines. The level of inhibition of phospho-p70S6 (T389), S6 (S235/236), GSK3B (S9), and PRAS40 (T246) was significant with copanlisib treatment in both H23AR and H358AR, similar to what was found in copanlisib-treated parental cells (H23 and H358) (**Figure 4H**). The expression of p4E-BP1 was found to be upregulated in both H23AR and H358AR as compared with their sensitive counterparts (**Figure 4H** & **Figure 5E-F**), and the degree of inhibition by copanlisib was more limited in H358AR vs H358 and H23AR vs H23 cells. Taken together, copanlisib sensitivity in acquired resistant cells is associated with the downregulation of various downstream pathway molecules.

### PI3K knockout via CRISPR-Cas9 restores sotorasib sensitivity in acquired resistant cells by dephosphorylating 4E-BP1

PI3K expression is upregulated in resistant cells, and its inhibition by copanlisib sensitizes these cells to treatment. To further validate the role of PI3K in resistance, we generated PI3K alpha-subunit knockout clones in both H23AR and H358AR cells using CRISPR-Cas9 technology and evaluated their sensitivity to sotorasib.

In H23AR cells, four PI3K-knockout clones (H23AR^*PI3K-KO-2-15*^, H23AR^*PI3K-KO-2-17*^, H23AR^*PI3K-KO-3-3*^, and H23AR^*PI3K-KO-3-12*^) were verified via PI3K sequencing and western blot analysis, confirming the complete loss of PI3K/p110α expression (**Figure 5A**). Cell viability assays showed that all PI3K-knockout clones exhibited significantly increased sensitivity to sotorasib compared to the parental H23AR cells (p=0.035, p=0.0002, p=0.09, & p=0.012, respectively), although the degree of sensitivity varied among clones (**Figure 5B**). Similarly, in H358AR cells, three PI3K-knockout clones (H358AR^*PI3K-KO-1-4*^, H358AR^*PI3K-KO-2-4*^, and H358AR^*PI3K-KO-2-7*^) were generated and confirmed by western blot analysis, showing no detectable PI3K/p110α expression (**Figure 5C**). These PI3K-knockout clones also exhibited significantly increased sensitivity to sotorasib compared to the resistant H358AR cells (Figure 5D), indicating that PI3K inhibition plays a direct role in restoring sotorasib sensitivity in resistant cells.

Among the H23AR knockout clones, H23AR^*PI3K-KO-2-17*^ showed the highest sensitivity to sotorasib, with a response comparable to the original resistant H23AR cells (p=0.0002). In contrast, H23AR^*PI3K-KO-3-3*^ exhibited the lowest sensitivity to sotorasib, though still significantly more sensitive than the original H23AR-resistant cells (**Figure 5B**). To further investigate the molecular basis of this differential sensitivity, we analyzed downstream signaling pathways in the most and least sensitive clones.

Like the effects of copanlisib on molecular signaling in H23AR cells (Figures 4B & H), robust downregulation of p-AKT (S473 & T308), pS6 (S235/236), pPRAS40 (T246), and p4E-BP1 (S65) was observed in the highly sensitive H23AR^*PI3K-KO-2-17*^ clone. Notably, copanlisib treatment did not further reduce the levels of these signaling molecules in this clone (**Figure 5E**). In contrast, in the least sensitive PI3K-knockout clone (H23AR^*PI3K-KO-3-3*^), similar levels of p-AKT (S473 & T308), pS6 (S235/236), and pPRAS40 (T246) downregulation were observed; however, p4E-BP1 (S65) remained elevated, resembling the levels found in H23AR cells suggesting a critical role of 4E-BP1 in sotorasib sensitivity (**Figure 5F**).

The expression of p4E-BP1 (S65) was significantly upregulated in H23AR compared to H23, the expression was completely depleted in the highly sensitive PI3K-knockout clone (H23AR^*PI3K-KO-2-17*^). Conversely, in the least sensitive PI3K-knockout clone (H23AR^*PI3K-KO-3-3*^), p4E-BP1 (S65) levels remained high, maintaining partial resistance. However, treatment of this least sensitive clone with copanlisib effectively reduced p4E-BP1 (S65) levels, along with significant inhibition of p-AKT (S473 & T308), pS6 (S235/236), and pPRAS40 (T246) (**Figure 5F**). Importantly, copanlisib treatment completely abolished p4E-BP1 (S65) expression in H23AR^*PI3K-KO-3-3*^, reducing it to levels comparable to those in the highly sensitive clone H23AR^*PI3K-KO-2-17*^. Taken together, these findings suggest that *PI3K* knockout or inhibition by copanlisib restores sotorasib sensitivity in resistant cells, primarily through the suppression of p4E-BP1.

### Phosphorylation of 4E-BP1, which is dependent on both mTORC1 and mTORC2 kinases, is associated with acquired resistance, and 4E-BP1 knockout via CRISPR-Cas9 restores sotorasib sensitivity

4E-BP1 is a negative regulator of protein translation that, in its non-phosphorylated state, binds to the translation initiation factor eIF4E, thereby inhibiting 5’-cap-dependent translation (16). We have shown that in sotorasib-resistant H23AR and H358AR cells, the 4E-BP1 and phosphorylated 4E-BP1 (p4E-BP1 at S65) were significantly upregulated compared to their parental counterparts. Treatment with copanlisib or PI3K knockout effectively suppressed p4E-BP1 in resistant cells (**Figure 5E & F**). Here, we compared the level of activated p4E-BP1 among PI3K-knockout clones and its correlation with sotorasib sensitivity.

Among four PI3K-knockout clones in H23AR cells clones (H23AR^*PI3K-KO-2-15*^, H23AR^*PI3K-KO-2-17*^, H23AR^*PI3K-KO-3-3*^, and H23AR^*PI3K-KO-3-12*^), sotorasib sensitivity was inversely correlated with their respective p4E-BP1 (S65) expression (**Figure 6A**). The most sensitive clone, H23AR^*PI3K-KO-2-17*^, exhibited almost the lowest level of p4E-BP1 expression, whereas the least sensitive clone, H23AR^*PI3K-KO-3-3*^, retained relatively higher levels of p4E-BP1. Notably, p-S6 was downregulated in all PI3K-knockout clones, but its inhibition did not correlate with sotorasib sensitivity (**Figure 6A**).

**Figure 6:**
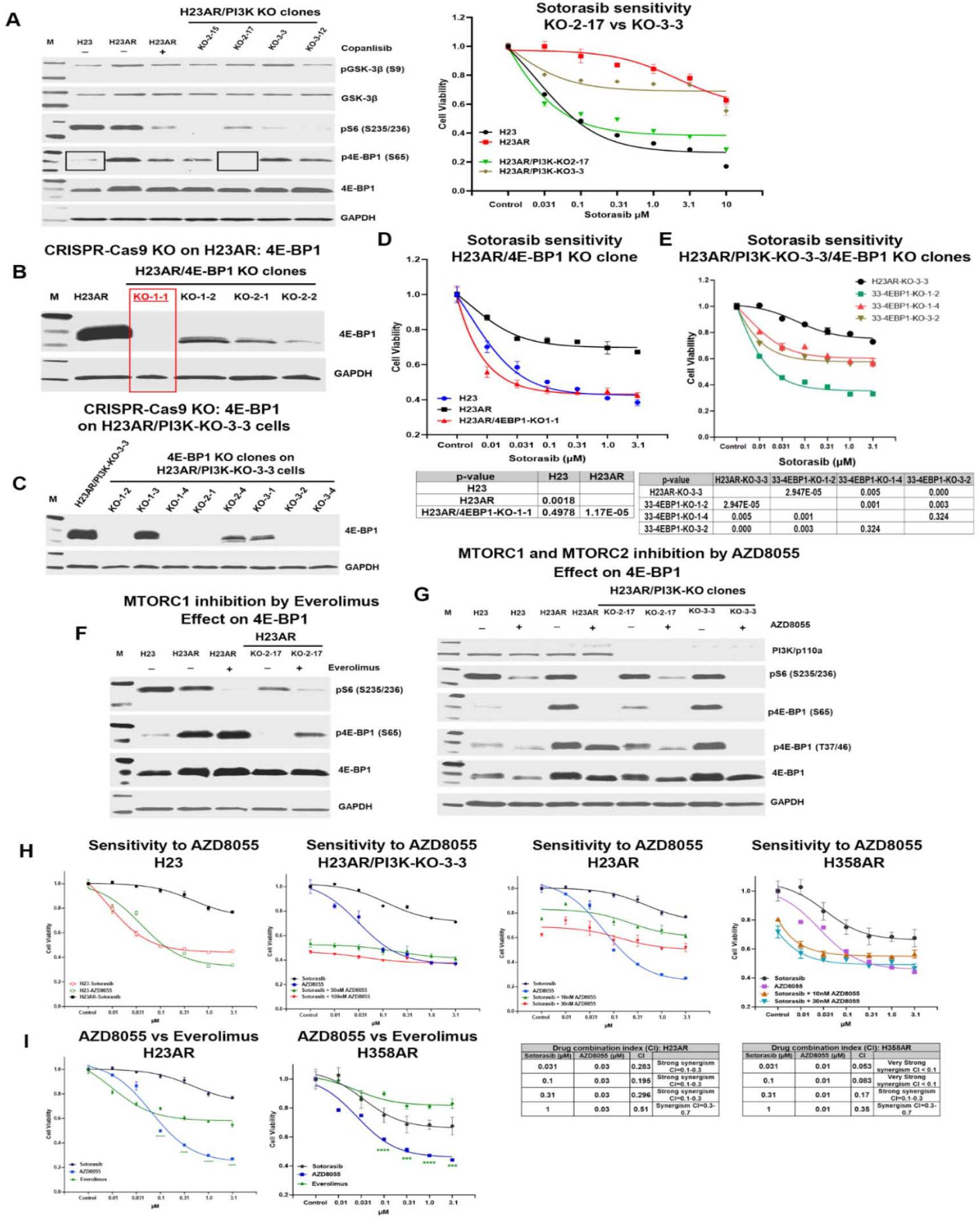
mTORC1/2 mediated phosphorylation of 4E-BP1 is associated with acquired resistance, and 4E-BP1 knockout via CRISPR-Cas9 restores sotorasib sensitivity. **A)** Association between 4E-BP1 expression and sotorasib sensitivity was determined among parental, acquired resistant, and PI3K knock-out clones. The level of expression of p-GSK-3B, p-S6, p-4E-BP1, and 4E-BP1 in H23 parental, H23AR resistant, and a series of PI3K knockout clones, H23AR^*PI3K-KO-2-15*^, H23AR^*PI3K-KO-2-17*^, H23AR^*PI3K-KO-3-3*^, H23AR^*PI3K-KO-3-12*^, was compared (left panel). The sotorasib sensitivity was compared among the most sensitive H23AR^*PI3K-KO-2-17*^, the least sensitive H23AR^*PI3K-KO-3-3*^ clones, and H23AR isogeneic cells (right panel). Higher expression of 4E-BP1 was associated with sotorasib sensitivity. Sotorasib sensitivity was assessed by treating the knockout and AR cells with varying drug concentrations for 72 hours and measuring cell viability using the Cell Glow assay. **B)** 4E-BP1 knockout clones were generated by the CRISPR-Cas9 technology on H23AR cells. The clones were confirmed based on the complete absence of 4E-BP1 expression in Western blots. H23AR^4E-BP1-KO-1-1^ clone was selected after screening several clones. **C)** Similarly, 4E-BP1 knockout clones were generated by CRISPR-Cas9 technology on sotorasib-least sensitive H23AR^*PI3K-KO-3-3*^ cells to make the double PI3K and 4E-BP1 knockout clones. Several clones were screened for their 4E-BP1 expression, and H23AR^*PI3K-KO-3-3/4E-BP1-KO-1-2*^, H23AR^*PI3K-KO-3-3/4E-BP1-KO-1-4*^, and H23AR^*PI3K-KO-3-3/4E-BP1-KO-3-4*^ clones were selected based on the complete deletion of 4E-BP1expression. **D)** Cell viability assay was performed for sotorasib sensitivity on 4E-BP1 knockout clone H23AR^*4E-BP1-KO-1-1*^ that was compared with parental H23 and H23AR cells. Statistical significance was determined based on p-value calculation, which is shown in the table below the graph. **E)** Cell viability assays were performed for sotorasib sensitivity on 4E-BP1 and PI3K double knockout clones H23AR^*PI3K-KO-3-3/4E-BP1-KO-1-2*^, H23AR^*PI3K-KO-3-3/4E-BP1-KO-1-4*^, H23AR^*PI3K-KO-3-3/4E-BP1-KO-3-4*^, and their sensitivity was compared with that of resistant H23AR cells. Statistical significance was determined based on p-value calculation, which is shown in the table below the graph. **F)** The effect of everolimus, an mTORC1 inhibitor, on the expression of p-S6 and p-4E-BP1 (S65) in resistant H23AR and sensitive PI3K knockout clone H23AR^*PI3K-KO-2-17*^ treated with everolimus for 24h. **G)** The effect of AZD8055, an mTORC1/2 dual inhibitor, on the expression of p-S6, p-4E-BP1 (S65), and p-4E-BP1 (T37/46) on parental H23, resistant H23AR, the most sensitive PI3K knockout clone H23AR^*PI3K-KO-2-17*^, and the least sensitive PI3K knockout clone H23AR^*PI3K-KO-3-3*^ after AZD8055 treatment for 24h. **H)** A cell viability assay was performed for AZD8055, mTORC1/2 dual inhibitor, and its combination with sotorasib on H23, H23AR, H358AR, and H23AR^*PI3K-KO-3-3*^ clone. Cell viability was measured using the Glow assay after 72 hrs of treatment at the designated doses. Drug synergy was calculated based on the combination index (CI) on both H23AR and H358AR cells. Combination Index (CI) is synergistic if the CI value is <0.7 (synergism if CI = 0.3-0.7, strong synergism if CI = 0.1-0.3, very strong synergism if CI <0.1). **I)** Cell viability assay on H23AR and H358AR cells to compare the cytotoxicity effect among Everolimus, AZD8055, and sotorasib. Each experiment was repeated three times. Data is shown as mean percentage ±SD, n ≥ 3.

To further examine the role of 4E-BP1 in resistance, we generated 4E-BP1-knockout clones using CRISPR-Cas9 in both H23AR cells and the least responsive PI3K-knockout clone (H23AR^*PI3K-KO-3-3*^). Western blot analysis confirmed the absence of 4E-BP1 expression in all selected clones, including one knockout clone in H23AR cells (H23AR^*4E-BP1-KO-1-1*^) and five clones in H23AR^*PI3K-KO-3-3*^ cells (H23AR^*PI3K-KO-3-3/4E-BP1-KO-1-2*^; H23AR^*PI3K-KO-3-3/4E-BP1-KO-1-4*^; H23AR^*PI3K-KO-3-3/4E-BP1-KO-2-1*^; H23AR^*PI3K-KO-3-3/4E-BP1-KO-3-2*^; H23AR^*PI3K-KO-3-3/4E-BP1-KO-3-4*^) (**Figure 6B & C**).

Cell viability assays showed that 4E-BP1 knockout significantly restored sotorasib sensitivity. The H23AR^*4E-BP1-KO-1-1*^ clone exhibited sensitivity comparable to parental H23 cells (p=0.00001 vs. H23AR; p=0.49 vs. H23) (**Figure 6D**).

All three 4E-BP1-knockout clones in the PI3K-KO cells (H23AR^*PI3K-KO-3-3/4E-BP1-KO-1-2*^; H23AR^*PI3K-KO-3-3/4E-BP1-KO-1-4*^; H23AR^*PI3K-KO-3-3/4E-BP1-KO-3-2*^) displayed significantly enhanced sotorasib sensitivity compared to their parental PI3K-knockout counterparts (p=0.00002, p=0.005, p=0.0001) (**Figure 6E**).

### mTORC1 and mTORC2 Mediate 4E-BP1 Hyperphosphorylation in Resistant Cells

Since p4E-BP1 (S65) was associated with sotorasib resistance and knocking down PI3K or 4E-BP1, or copanlisib treatment restores sotorasib sensitivity in acquired resistant cells, we further investigated the role of mTORC1 and mTORC2 in 4E-BP1 phosphorylation in resistant cells. Everolimus, a mTORC1 inhibitor, and AZD8055, a dual mTORC1/2 inhibitor, were used to treat H23 parental, H23AR, H23AR^*PI3K-KO-2-17*^ (most sensitive clone), and H23AR^*PI3K-KO-3-3*^ (least sensitive clone) cells.

Everolimus treatment significantly downregulated pS6 (S235/236) in H23AR and H23AR^*PI3K-KO-2-17*^, but did not reduce p4E-BP1 (S65) levels (Figure 6F). Everolimus induced only moderate cytotoxicity in resistant cells and organoids, as observed in cell viability assays (**Figure 4I and 6I**). These findings suggest that while mTORC1 inhibition suppresses pS6 phosphorylation, it does not directly regulate p4E-BP1 in resistant cells, implicating a mTORC1-independent kinase in 4E-BP1 phosphorylation.

In contrast, treatment with AZD8055, a dual mTORC1/2 inhibitor, completely abolished p4E-BP1 (S65) expression in both parental H23 and resistant H23AR cells, as well as in both the least and most sensitive PI3K-knockout clones (H23AR^*PI3K-KO-3-3*^ and H23AR^*PI3K-KO-2-17*^, respectively) (**Figure 6G**). pS6 downregulation was observed in all cells, irrespective of their sotorasib sensitivity.

### mTORC1/2 Dual Inhibition Restores Sotorasib Sensitivity in Resistant Cells

Dual inhibitor AZD8055 treatment significantly reduced cell viability in sotorasib-resistant H23AR, H358AR, and the least sensitive PI3K-knockout clone (H23AR^*PI3K-KO-3-3*^) (**Figure 6H**). When comparing AZD8055 with everolimus in resistant H23AR and H358AR cells, the dual inhibitor induced significantly greater cytotoxicity than the mTORC1-specific inhibitor (**Figure 6I**).

Further analysis of the combination index (CI) between sotorasib and AZD8055 revealed strong synergy in resistant cells. In H23AR cells, sotorasib (0.031 µM) combined with AZD8055 (0.03 µM) yielded a CI of 0.283, indicating strong synergism. In H358AR cells, sotorasib (0.031 µM) combined with AZD8055 (0.01 µM) exhibited an even stronger synergy, with a CI of 0.053 (**Figure 6H**).

This data suggests that dephosphorylation of 4E-BP1 through PI3K knockout, 4E-BP1 knockout, or copanlisib treatment restores sotorasib sensitivity. Notably, mTORC1 inhibition alone is insufficient to suppress p4E-BP1, while dual inhibition of mTORC1 and mTORC2 effectively abolishes p4E-BP1 expression, leading to robust cytotoxicity and enhanced sotorasib sensitivity indicating the suppression of p4E-BP1 is dependent on both mTORC1 and mTORC2 complexes.

### Overcoming Resistance by Targeting PI3K-AKT-mTOR Signaling with copanlisib or sapanisertib in acquired resistant NSCLC PDX, CDX, and Xenograft Tumors

Our findings thus far indicate that PI3K inhibition, either through copanlisib treatment or genetic knockout, restores sotorasib sensitivity in resistant cells by suppressing p4E-BP1, a process mediated by both mTORC1 and mTORC2 kinases. We previously showed that the dual mTORC1/2 inhibitor, AZD8055, enhances sotorasib sensitivity and synergistically inhibits cell proliferation in resistant models. To further evaluate the therapeutic potential of these inhibitors, we examined their efficacy in vivo in sotorasib-resistant patient-derived xenografts (PDX), cell-derived xenografts (CDX), and other xenograft models.

Cell viability assays confirmed that copanlisib was cytotoxic for sotorasib-acquired resistant H23AR and H358AR cells, as well as primary and acquired sotorasib-resistant organoids (PDXOs) (**Figure 4A & C**). Furthermore, copanlisib not only restored sotorasib sensitivity in H23AR and H358AR cells but also exhibited strong synergy when combined with sotorasib (**Figure 7A**).

**Figure 7:**
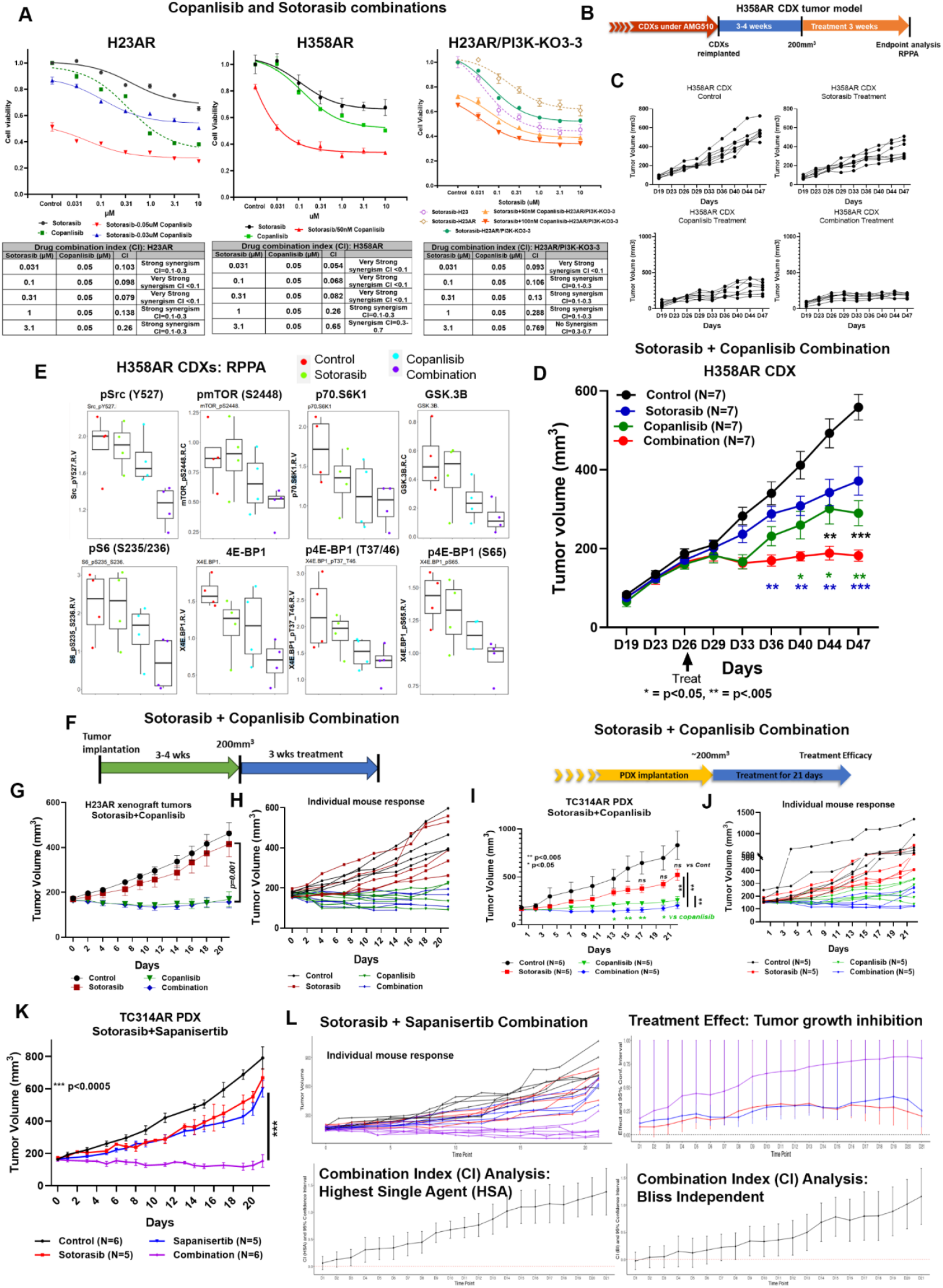
Overcoming resistance by targeting PI3K-AKT-mTOR signaling with copanlisib or sapanisertib in acquired resistant NSCLC PDX, CDX, and xenograft Tumors. **A)** In vitro cell viability assay was performed to evaluate the synergistic effect of sotorasib + copanlisib on H23AR, H358AR, and H23AR^PI3K-KO-3-3^ cells. Cells were treated with sotorasib, copanlisib, and their combination. In combination, very low fixed copanlisib doses (either 0.05µM or 0.03µM) were combined with variable doses of sotorasib and cell viability assays were performed after 72h of treatment. The combination index (CI) was calculated and presented in the table listed under the graph. The Combination Index (CI) is synergistic if the CI value is <0.7 (synergism if CI = 0.3-0.7, strong synergism if CI = 0.1-0.3, very strong synergism if CI <0.1). **B-D)** In vivo antitumor efficacy of copanlisib and sotorasib combination on acquired resistant H358AR CDX tumors. Mice were engrafted with freshly harvested 3^rd^-generation acquired-resistant H358AR CDX tumor tissues into experimental mice. When the tumor volumes reached approximately 200 mm^3^, mice were randomized into four treatment groups (control, sotorasib (100 mg/kg), copanlisib (6 mg/kg), and sotorasib + copanlisib combination) and treated for at least 3 weeks. **B)** Treatment strategy is shown, **C)** Individual mouse response for each group. **D)** Antitumor effect of copanlisib and sotorasib combination on H358AR CDX tumors. **E)** The residual tumor tissues from H358AR CDX tumors were harvested and subjected to RPPA analysis. The level of p-Src, p-mTOR, p70.S6, GSK-3b, p-S6, 4E-BP1, p-4E-BP1 (T37/46), and p-4E-BP1 (S65) expression in residual tissues was determined and compared among the treatment groups. **F-H)** Antitumor effect of sotorasib + copanlisib combination on acquired resistant H23AR xenograft tumors. **F)** The treatment strategy is shown, where 5-8 million cells/mouse were subcutaneously injected. When tumor sizes reached 200 mm^3^, tumors were treated for 3 weeks with sotorasib (50 mg/kg), copanlisib (6 mg/kg), and the combination. **G)** Antitumor effect is shown. **H)** Individual mouse responses were compared with mice from every treatment group. **I)** Antitumor effect of sotorasib + copanlisib combination on acquired resistant TC314AR PDX tumors. Mice were engrafted with freshly harvested 3^rd^-generation acquired resistant TC314AR PDX tumor tissues into experimental mice. When the tumor volumes reached approximately 200 mm^3^, mice were randomized into four treatment groups (control, sotorasib (50 mg/kg), copanlisib (6 mg/kg), and sotorasib+copanlisib combination) and treated for 3 weeks. Tumor growth curves were generated for each treatment group from the beginning to the end of the treatment. **J)** Individual mouse responses were compared with mice from every treatment group. **K)** Antitumor effect of sotorasib + sapanisertib combination on acquired resistant TC314AR PDX tumors. Mice were engrafted with freshly harvested 3^rd^-generation acquired resistant TC314AR PDX tumor tissues into experimental mice. When the tumor volumes reached approximately 200 mm^3^, mice were randomized into four treatment groups (control, sotorasib (50 mg/kg), sapanisertib (1 mg/kg), and sotorasib + sapanisertib combination) and treated for 3 weeks. Tumor growth curves were generated for each treatment group from the beginning to the end of the treatment. **L)** The antitumor data were analyzed by a statistical program, CombPDX, to evaluate the synergistic antitumor effect between sotorasib and sapanisertib on this TC314AR PDX model. Top left) shows the individual mouse response. Top right) Treatment effect for combination, sotorasib alone, and sapanisertib alone treatment. The treatment effect was calculated based on the percentage of tumor growth inhibition with a 95% confidence interval. Bottom panels) The combination index (CI) was calculated by using two different methods, including Highest Single Agent analysis (HAS) and Bliss Independent (BI) analysis with a 95% confidence interval. In-vivo experiments were repeated three times with 5-7 mice/group used in each experiment. Statistics are shown at a significance level of p<0.05 unless otherwise noted. Data is shown as mean percentage ±SD, n=5. *, *P* < 0.05; **, *P* <0.005; *** *P* <0.0005.

Combination index (CI) analysis revealed a strong to very strong synergism when sotorasib (0.031– 0.31 µM) was combined with copanlisib (0.05 µM) in both H23AR and H358AR cells (**Figure 7A**). Notably, the least responsive PI3K-knockout clone (H23AR^*PI3K-KO-3-3*^) also exhibited significant sensitivity to the combination treatment, with a CI of 0.093 at sotorasib (0.031 µM) + copanlisib (0.05 µM) (**Figure 7A**).

To evaluate the in vivo efficacy of this combination, we established sotorasib-resistant H358AR-CDX tumors in NSG mice. Once tumor volumes reached approximately 200 mm^3^, mice were treated with copanlisib, sotorasib, or their combination, following the strategy outlined in **Figure 7B**.

Copanlisib monotherapy showed significant antitumor activity compared to the control group. However, the combination of sotorasib and copanlisib exhibited a synergistic antitumor effect, with statistically significant tumor suppression observed at D40, D44, and D47 (**Figure 7D**). The combination treatment was superior to either monotherapy in controlling tumor growth.

Individual tumor growth curves (**Figure 7C**) revealed that in the combination group, tumor volumes remained stable, with no significant growth observed between pre-treatment (D26) and post-treatment (D44) measurements. In contrast, tumors in the sotorasib monotherapy group continued to grow significantly between D26 and D44, further highlighting the superior efficacy of the combination strategy.

To investigate the underlying mechanism, tumors were harvested after three weeks of treatment and analyzed using RPPA. The expression levels of total 4E-BP1, phosphorylated 4E-BP1 at T37/46 and S65 were significantly suppressed by both copanlisib monotherapy and the combination treatment (**Figure 7E**). Additionally, other key downstream effectors of the PI3K-AKT-mTOR pathway, including phosphorylated S6 (S235/236), phosphorylated mTOR (S2448), GSK-3β, phosphorylated Src (Y527), and p70-S6K1, were also downregulated (**Figure 7E**). In contrast, sotorasib monotherapy failed to reduce the phosphorylation of key signaling proteins such as p-mTOR (S2448), p-S6 (S235/236), p4E-BP1 (S65), and p-Src (Y527). This indicates that the PI3K-AKT-mTOR signaling pathway remained functionally active in resistant tumors, thereby sustaining resistance to sotorasib treatment.

The combination of sotorasib and copanlisib was evaluated to determine its in vivo efficacy in overcoming sotorasib-acquired resistance in H23AR xenograft tumors. H23AR cells were continuously cultured under sotorasib pressure during cell expansion to develop resistance, and these resistant cells were subsequently implanted into NSG mice. Treatment was initiated once the tumor volumes reached approximately 200 mm^3^, following the experimental strategy illustrated in **Figure 7F**. When administered as a monotherapy, sotorasib did not demonstrate a statistically significant antitumor effect against H23AR xenografts compared to the control group (p = 0.40). However, treatment with copanlisib alone resulted in a strong antitumor response, which was statistically significant when compared to both sotorasib monotherapy and the control group (p < 0.001) (**Figure 7G**). Although the combination of copanlisib and sotorasib did not exhibit a synergistic effect due to the strong copanlisib single agent effect (p = 0.82; comb vs copanlisib), the combination therapy successfully overcame sotorasib resistance by significantly inhibiting tumor growth compared to both sotorasib monotherapy (p = 0.001) and the control group (p < 0.001) (**Figure 7G**). Individual mouse responses showed that tumor growth was inhibited in every mouse treated with the combination treatment (**Figure 7H**).

Like the H23AR xenograft model, the therapeutic efficacy of sotorasib in combination with copanlisib was also investigated in the TC314AR patient-derived xenograft (PDX) model, which has acquired resistance to sotorasib. TC314AR-resistant PDXs were implanted into NSG mice, and the experimental design followed an individualized PDX trial approach. In this strategy, each mouse was blindly assigned to one of the four treatment groups—control, sotorasib, copanlisib, or combination once the PDX tumor volume reached approximately 200 mm^3^. The treatment regimen was continued for three weeks to assess the antitumor effects of each therapy. As anticipated, sotorasib monotherapy failed to exert a significant inhibitory effect on tumor growth when compared to the control/vehicle-treated group (p = ns vs control). In contrast, copanlisib monotherapy showed a potent antitumor response, which was statistically significant when compared to both the control and sotorasib monotherapy groups (p <0.005). The combination of sotorasib and copanlisib produced an even stronger tumor-suppressive effect than either monotherapy (**Figure 7I**). Individual mouse responses are shown in **Figure 7J**. The efficacy of this combination was analyzed using the CombPDX tool, a unified statistical platform designed to evaluate drug synergy in PDX models (17). The treatment effect shown in **Figure 7-figure Supplement 1A** was measured based on tumor growth inhibition (TGI) with a 95% confidence interval in comparison with the control group, and the combination treatment showed superior efficacy. The Combination Index (CI), calculated based on the Highest Single Agent (HSA) method, yielded a positive value, indicating a synergistic interaction between sotorasib and copanlisib (**Figure 7-figure supplement 1B**).

We showed above that the dual mTORC1/mTORC2 inhibitor AZD8055 was synergistic in vitro against sotorasib-resistant cells and PDX organoids (**Figure 6H**). We therefore investigated the antitumor efficacy of combining sotorasib with sapanisertib, another dual mTORC1/mTORC2 inhibitor, in the TC314AR PDX model. In addition to AZD8055, another dual mTORC1/mTORC2 inhibitor, sapanisertib, was tested in vitro and in vivo because the drug is currently used in the clinic for many cancer types. In vitro cell viability assays showed that sapanisertib exhibited a cytotoxic effect, significantly inhibiting the proliferation of acquired-resistant cell lines, including H23AR, H358AR, and PDXO models (**Figure 7-figure Supplement 2A-B**). TC314AR PDX tumors were established in NSG mice, and once tumor volumes reached approximately 200 mm^3^, treatment with either sotorasib monotherapy, sapanisertib monotherapy, or the combination of both agents was administered for three weeks. As observed in prior resistance models, sotorasib alone did not elicit a significant antitumor response, with no notable tumor growth inhibition observed. In contrast, the combination of sotorasib with sapanisertib led to a dramatic and highly significant antitumor effect (**Figure 7K**). This combination therapy was further evaluated using the CombPDX analysis tool (17), a comprehensive statistical tool to analyze the drug synergy on PDXs, which confirmed that the antitumor response observed with the combination was markedly superior to that of either monotherapy. Tumor growth inhibition (TGI) was assessed with a 95% confidence interval, revealing that the combination treatment resulted in a TGI value exceeding 0.75 (on a scale of 0 to 1.0), whereas sotorasib and sapanisertib monotherapies showed TGI values of less than 0.2 (**Figure 7L**). The Combination Index (CI) was calculated as previously reported and indicated synergy for the combination (23). (**Figure 7L**).

These findings underscore the critical role of PI3K-AKT-mTOR signaling in sotorasib resistance and show that targeting this pathway with copanlisib or dual mTORC1/2 inhibition can effectively restore sensitivity to sotorasib. The combination of sotorasib with copanlisib exhibited robust synergy both in vitro and in vivo, providing a promising therapeutic strategy to overcome acquired resistance in PDX and CDX tumor models.

## Discussion

Due to the intrinsic characteristics of KRAS proteins, KRAS has been considered an undruggable target. The recent advent of compounds allowing direct inhibition of the *KRAS*^*G12C*^ oncogene, such as adagrasib and sotorasib, has initiated a new era in the clinical management of *KRAS*^*G12C*^ mutant NSCLC. Both AMG510 (Sotorasib) and MRTX849 (Adagrasib) bind covalently to the GDP-bound form of the *KRAS*^*G12C*^ protein (KRAS OFF), preventing oncogenic signaling and causing tumor regression in preclinical models (9,18,19). Current *KRAS*^*G12C*^ (OFF) inhibitors cause responses in less than half of patients, and these responses are not durable. Other than sotorasib and adagrasib, other new inhibitors are being developed, targeting either the KRAS active state (KRAS ON), where RAS is coupled with GTP, or inactive state (KRAS OFF), where RAS binds with GDP, and evaluating the clinical efficacy (20). In early-phase clinical trials, adagrasib and sotorasib have shown promising results against non–small cell lung cancer and more modest efficacy against colorectal cancer (9,11). Despite high initial efficacy, there is a relatively short progression-free survival due to resistance that develops in virtually all patients to single-agent therapy. Moreover, the responsiveness of NSCLC to *KRAS*^*G12C*^ inhibitors is not sufficient compared with that of TKIs such as EGFR-TKI and ALK-TKI, which show response rates of 70% and 80%, respectively (21,22).

An improved understanding of the biological basis of acquired resistance is necessary, which will provide more opportunities to optimize *KRAS*^*G12C*^ inhibitor regimens and new combinations. Multiple mechanisms of resistance have been described in the clinic upon treatment with direct *KRAS* inhibitors (23,24). Acquired resistance to KRAS^G12C^ inhibitors is mainly driven by genomic mutations and non-genetic alteration events (25). Development of secondary mutations in *KRAS*^*G12C*^ oncoprotein that interfere with the binding of covalent inhibitors to cysteine 12 residues, or mutations in the normal *KRAS* alleles that could not be targeted by the KRAS inhibitors, or other mutations in BRAF, MAP2K1, RTKs that are related to the reactivation of the RAS-RAF-MEK-ERK signaling pathway are the major genetic events associated with sotorasib resistance (26,27). Other tumors displayed mutations in *PI3K, PTEN, SMAARCA4, KEAP1*, and *PTCH1* effectors in additional oncogenic pathways (26,28-30). In this study, sotorasib acquired resistant isogenic patient-derived xenografts (PDXs: TC314AR and TC303AR), CDX, PDX-derived organoids (PDXOs: (PDXO314AR & PDXO303AR), and H23AR and H358AR cell lines were developed and investigated to explore the underlying mechanism of resistance. Whole exome sequence was performed in all acquired resistant models; no additional mutations were found in *KRAS*, and no new mutations were found in major proteins associated with bypass pathways. Although each resistant model has its distinct molecular profiling pattern, all resistant models retained the *KRAS*^*G12C*^ mutation regardless.

Yet about half of the resistant tumors examined did not display additional mutations, thus indicating the existence of additional mechanisms that might cause resistance to KRAS inhibition (23,31). Identification of these resistance mechanisms is an urgent prerequisite to developing improved therapeutic strategies. Reactivation of KRAS and/or upregulation of additional bypass signaling pathways are found to be one of the potential central mechanisms of KRASi-related acquired resistance. Reactivation of the MAPK pathway without developing new mutations in *KRAS* or its downstream mediators has been reported in *KRAS*^*G12C*^ lung adenocarcinomas resistant to the KRAS^G12C^ inhibitor AMG 510 and adagrasib (27,32). Consistent with that, we also found upregulation of ERK signaling determined by the level of phosphorylation of ERK1/2 in sotorasib-resistant H23AR and H358AR cells, which was inhibited by the treatment of KRASi. On the other hand, the complete inhibition of ERK1/2 phosphorylation by sotorasib treatment was noticed in parental H23 and H358 cells. This hyper-phosphorylation of EKR1/2 in acquired resistant cells could be explained by the hyper-formation of new KRAS^G12C^ proteins and reactivation of downstream signaling in resistant cells (33). High-throughput Mass-Spectrometry (MS) was performed on acquired resistant TC314AR and TC303AR PDXs to understand the bypass resistance mechanisms. Enrichment analysis showed that PI3K-AKT-mTOR signaling was significantly upregulated in resistant PDXs as compared with their parental counterpart. Over-expression of PI3K was also found in acquired resistant cell lines, which was found in consistent with other studies where activation of the PI3K-AKT-mTOR canonical pathway was found as an additional bypass mechanism in acquired resistance (23,24). Other alternative pathways, such as epidermal growth factor receptor or Aurora kinase A (AURKA) signaling, FAK, and Hippo pathways, are also reported to be associated with resistance to KRAS inhibitors (23,34). These alternative pathways were not found to be significantly upregulated in our study, indicating heterogeneous mechanisms involved in KRASG12C inhibitor resistance, notably being tissue-specific.

The PI3K–AKT–mTOR signaling pathway is frequently activated in cancer, deregulating control of metabolism, cell proliferation, and apoptosis (35). Pharmaceutical inhibition of PI3K by copanlisib or genetic knock-out of PI3K by CRISPR-Cas9 restores the sensitivity of sotorasib in sotorasib-resistant cells or organoids. Copanlisib is a potent pan-PI3K inhibitor, an FDA-approved compound that has been shown to inhibit the cellular proliferation of relapsed or refractory, indolent, or aggressive lymphoma (36). However, there are no studies investigating the use of copanlisib in NSCLC preclinical models. PI3K expression was upregulated in sotorasib-resistant H23AR and H358AR cells, and inhibition of PI3K by copanlisib induced cytotoxicity in the cells. Similarly, sotorasib-acquired resistant PDXOs such as PDXO314AR and PDXO303AR were also found to be very sensitive to copanlisib.

We have developed around PDXOs from their respective PDXs. Over 50% of those PDXOs were primarily resistant to sotorasib or adagrasib. No additional KRAS mutation was found in any of those PDXs, and no significant correlation was found with any co-mutation with either PI3K, STK11, KEAP1, SMARCA4, etc. However, no co-mutation was associated with sotorasib resistance. As a majority of PDXOs showed primary resistance to sotorasib, this suggests that other bypass signaling mechanisms are involved in sotorasib resistance. Primarily resistant PDXOs were found to be very sensitive to copanlisib in the PDXO viability assay, suggesting the role of PI3K-AKT-mTOR signaling in the inactivation of sotorasib sensitivity. Copanlisib overcame sotorasib resistance in the sotorasib-resistant H23AR and H358AR cells and PDXOs (PDXO303AR and PDXO314AR) and worked synergistically with sotorasib in-vitro and in-vivo. The drug combination index (CI) analysis showed that a very low dose of copanlisib (50nM) worked synergistically in sensitizing resistant cells to sotorasib, which could mitigate copanlisib-associated drug toxicity.

To understand the underlying resistance mechanism of the PI3K-AKT-mTOR pathway, blocking the upstream PI3K by pan-PI3K inhibitor copanlisib, which inhibits the catalytic activity of the class I PI3Kα, β, γ, and δ isoforms, inhibited AKT phosphorylation at both T308 and S347 and its direct substrates PRAS40 (T246) and GSK3β (S9), which was consistent with breast tumor models (35,37). Inhibition of AKT by MK2206, an AKT inhibitor, moderately inhibited the colony formation and worked significantly better when combined with copanlisib, suggesting that overcoming resistance by copanlisib is not solely dependent on AKT inhibition. The moderate activity in resistant cells by AKT inhibitor, MK2206, can be explained by its moderate inhibition of downstream mTORC1 targets. Will et al reported that targets of mTOR, such as pS6 and p4EBP1, were also inhibited by both MK2206 and copanlisib, although PI3K inhibition blocked 4E-BP1 phosphorylation more potently and longer than AKT inhibition (35). In our study, copanlisib was found to be very effective in downregulating the mTORC1 signaling through inhibiting the phosphorylation of the target molecules such as S6 and 4E-BP1.

Like AKT inhibition by MK2206, inhibition of mTORC1 by everolimus, which targets S6 and inhibits the phosphorylation of S6, inhibited colony formation moderately in sotorasib-resistant H23AR and H358AR cells, but the inhibition of colony formation increased significantly when combined with copanlisib. In cell viability assays in acquired resistant isogeneic cells or PDXOs, copanlisib was found to be very effective, and the effect was much stronger for copanlisib compared to everolimus. This data suggested that the downstream mechanism of PI3K inhibition by copanlisib involves more than S6-mediated mTORC1 activity. To verify the precise involvement of mTORC1, we investigated two downstream effector molecules, S6 and 4E-BP1, which are involved in protein translation. Both S6 and 4E-BP1 phosphorylation were significantly inhibited by copanlisib treatment in resistant cells, although basal expression of 4E-BP1 was significantly higher only in resistant cells. Everolimus, an mTORC1 inhibitor, downregulated S6 phosphorylation in resistant cells where 4E-BP1 expression remained unaffected, indicating dephosphorylation of 4E-BP1 is essential for overcoming sotorasib resistance.

4E-BP1 is a member of the 4E-BP family that represses translation by competing with eIF4G for binding to eIF4E, thereby preventing the formation of the eIF4F complex. In cancer cells, however, 4E-BP1 is frequently phosphorylated by its upstream oncogenic signals such as PI3K/AKT and RAS/RAF/MEK/ERK pathways, which causes 4E-BP1 disassociation from eIF4E and increases cap-dependent protein translation. The mTOR kinase complex 1 (mTORC1) is the master regulator of eIF4E-initiated cap-dependent translation by phosphorylating 4E-BP1 on Thr37 and Thr46 which act as priming sites for its subsequent phosphorylation on Ser65 (38). Hyperphosphorylation of 4E-BP1 was found in sotorasib-resistant cells as compared with sensitive cells, which was consistent with other studies that found an association with malignant progression and poor prognosis with 4E-BP1 hyperphosphorylation (39,40). PI3K inhibition by copanlisib sensitizes the cells by downregulating 4E-BP1, which emphasizes the role of 4E-BP1 in overcoming sotorasib resistance. We also found a clear inverse association between the level of sotorasib cytotoxicity and phosphorylation of 4E-BP1 among PI3K-knockout clones. The clone H23AR-PI3K-KO-2-17, the most sensitive clone, showed complete inhibition of phosphorylation of 4E-BP1 (S65), which resembles the status found in the parental H23 cells. On the other hand, the clone H23AR-PI3K-KO-3-3, the least sensitive clone, showed partial dephosphorylation of 4E-BP1 and correlated with the expression found in sotorasib-resistant H23AR cells. To verify the precise role of 4E-BP1, CRISPR-Cas9 knock-out of 4E-BP1 clones was created in resistant cells. After knocking out 4E-BP1, both resistant cells, H23AR and H358AR, became sensitive to sotorasib. When 4E-BP1 was knocked out from sotorasib’s least sensitive PI3K knock-clone (H23AR-PI3K-KO-3-3), the double knock-out clone (PI3K and 4E-BP1) restored sotorasib sensitivity to the level of sensitive cells, suggesting the pivotal role of 4E-BP1 in overcoming sotorasib resistance.

4E-BP1 is the downstream target of mTORC1. To verify the dependency on mTORC1 kinase on 4E-BP1 phosphorylation and subsequent protein translation, everolimus (mTORC1-specific inhibitor) and AZD8055, a dual inhibitor that inhibits both mTORC1 and mTORC2, were used to treat the resistant cells. Everolimus can stop S6 phosphorylation without having any effect on 4E-BP1 in resistant cells, whereas AZD8055 reduced p4E-BP1 and pS6 in resistant cells, suggesting the hyperphosphorylation of 4E-BP1 in resistant cells was independent of mTORC1. ADZ8055 dual mTOR inhibitor, also synergized with sotorasib in restoring the sensitivity in sotorasib-resistant cells and PDXOs. In our study, mTORC1 inhibition by everolimus did not restore sotorasib (KRAS OFF inhibitor) sensitivity, but a recent study found a synergistic effect of KRASi (KRAS ON inhibitor) with mTORC1 kinase inhibition (41). Another recent study showed that RAS-ON inhibition can overcome resistance developed from RAS-OFF blockade, which is sotorasib or adagrasib (42). In conclusion, our study found a strong synergistic antitumor effect of sotorasib with upstream PI3K inhibitor or downstream mTORC1 and mTORC2 dual inhibitor in sotorasib-acquired resistant PDXs, CDX, and xenograft tumors, suggesting a possible combination to overcome the acquired resistance that merits translation into clinical trials.

## Materials and Methods

### Sotorasib-acquired resistant cell lines, cell culture, and maintenance

The human parental NCI-H23 and NCI-H358 NSCLC cell lines, which carry *KRAS*^*G12C*^ mutations, were obtained from ATCC, and their sotorasib-resistant isogenic clones, H23AR and H358AR, were obtained from Dr. Srikumar P. Chellappan’s laboratory (Moffitt Cancer Center, Tampa, FL). H23 and H358 cells were cultured and maintained in RPMI-1640 complete media supplemented with 10% heat-inactivated fetal bovine serum (GE Healthcare Life Sciences, HyClone Laboratories) and 1% penicillin-streptomycin (Thermo Fisher Scientific). The H23AR and H358AR resistant clones were cultured in RPMI-1640 complete medium with 2.5 µM and 1 µM AMG510 (Sotorasib) (Medchemexpress (MCE), NJ, USA), respectively, which was dissolved in DMSO, stored at -70°C, and diluted in culture medium for in-vitro experiments. *KRAS*^*G12G*^ inhibitors, including sotorasib, adagrasib, and opnurasib, were purchased from MedChemExpress (MCE), NJ, USA. Other targeted molecules used in this study, such as copanlisib (PI3K inhibitor), AZD8055 (mTORC1/2 inhibitor), Everolimus (mTORC1 inhibitor), MK2206 (AKT inhibitor), and Sapanisertib (mTORC1/2 inhibitor), were purchased from Selleckchem (Houston, TX, USA). All cell lines tested negative for mycoplasma before use in experiments. The cell lines were tested for mycoplasma routinely by ELISA in a core lab at MD Anderson Cancer Center. The cell lines were also authenticated before the experiment by the core lab at The University of Texas MD Anderson Cancer Center.

### In-vivo animal studies

NOD. Cg-*Prkdc*^*scid*^ *Il2rg*^*tm1Wjl*^/SzJ (NSG) mice were obtained from The Jackson Laboratory, which were used for subcutaneous patient-derived xenograft (PDX), Cell line-derived xenograft (CDX), and xenograft tumors development for sotorasib acquired resistant studies. Female 6-to-8-week-old mice were used in these studies. Mice were housed in micro-isolator cages under specific pathogen-free conditions in a dedicated mice room in the animal facility at The University of Texas MD Anderson Cancer Center. Mice were given autoclaved acidified water and fed a special diet (Uniprim diet). All animal experiments were carried out under approved IACUC (Institutional Animal Care and Use Committee (IACUC) protocols following approval by the MDACC institutional review board, and were performed by the Guidelines for the Care and Use of Laboratory Animals published by the National Institutes of Health.

H23 and H23AR xenograft tumors were generated by subcutaneously injecting 5–8 million cells per mouse into NSG mice. To ensure the maintenance of acquired resistance, H23AR cells were expanded in cell culture with sotorasib (2.5 µM) before implantation. Tumors were allowed to develop over a period of 4–6 weeks, after which all tumor-bearing mice were randomized blindly into different treatment groups, as specified in each in vivo experiment.

Once tumor volumes reached approximately 200 mm^3^, treatment was initiated and continued for at least three weeks. In studies evaluating the combination of sotorasib and copanlisib, single-agent treatment groups were included, with each group consisting of at least 5–7 mice. Control group mice received vehicle treatment, while the experimental groups were treated with sotorasib (50 mg/kg) and copanlisib (6 mg/kg), administered once daily (QD) for five days per week over a three-week period.

Mice were monitored daily for potential side effects, and tumor growth was assessed by measuring two perpendicular tumor diameters two to three times per week. Tumor surface area was calculated using the formula: 1/2 × (Length × Width^2^).

### Development of sotorasib-acquired resistant cell line-derived xenograft (CDX) model

The *KRAS*^*G12C*^-mutant H358 cell line, recognized as one of the most sensitive models to sotorasib treatment, was implanted into NSG mice via subcutaneous injection, with each mouse receiving approximately 5–8 million cells. Once the tumors reached a volume of 200 mm^3^, the mice were administered sotorasib (AMG510) at a dosage of 100 mg/kg once daily (QD) for five days per week over a period of two months.

Tumors that grew beyond 500 mm^3^ were harvested, reimplanted into new NSG mice, and allowed to establish at 200 mm^3^ before initiating a second round of sotorasib treatment for an additional two months. Following this treatment period, the first-generation (G1) resistant tumors were collected and reimplanted into a new cohort of mice, where they were subjected to another four months of sotorasib treatment to generate the second-generation resistant CDX (G2). These G2-resistant CDXs were subsequently reimplanted into new mice and underwent another two-month cycle of sotorasib treatment to develop the third generation (G3) H358AR CDX model.

To evaluate the resistance potential of the G3 CDX model, G3 H358AR CDXs were implanted into a fresh batch of NSG mice, while a separate group of NSG mice received subcutaneous injections of parental H358 cells. Once tumors from both the G3 H358AR CDX and H358 xenografts reached 200 mm^3^, sotorasib treatment was initiated and maintained for three weeks. Tumor growth inhibition was assessed at Day 21 of treatment, revealing that the G3 H358AR CDX exhibited no detectable antitumor response. Whole-exome sequencing (WES) was performed on these CDXs to confirm the presence of the *KRAS*^*G12C*^ mutation.

### Development and therapeutic studies on NSCLC PDX models

12 *KRAS*^*G12C*^ mutant PDXs were selected from our core with over 200 NSCLC PDXs in UT MD Anderson Cancer Center (Dr. Jack Roth, PI) developed under the NCI–funded PDX Development and Trial Centers Research Network (PDXNet) program. All the PDXs were well characterized and well annotated based on RNASeq, whole exome sequencing (WES) (43). PDXs were established from frozen stocks through implantation into NSG mice according to the protocol published previously (44,45). The PDX studies were conducted according to the approved institutional animal care and use committee protocol in accordance with the procedures outlined in the Guide for Care and Use of Laboratory Animals.

#### For the sotorasib sensitive study

PDXs TC247, TC314, TC303, and TC453 were established in NSG mice simultaneously or 1-2 models at a time. Tumor-bearing mice were randomized to treatment and control groups (n=5-7 mice/group) when tumor volume reached approximately 200 mm^3^. Treatments were started simultaneously for the models within an acceptable size window (such as 200 mm^3^) or independently for each mouse when a certain tumor size was reached (e.g., at a predetermined size such as 200 mm^3^) if mice were enrolled in all groups in a parallel fashion. PDX-bearing mice were treated with sotorasib at a dose of 100 mg/kg or 50 mg/kg unless otherwise mentioned. The treatment was continued for 5 days/week for 3 weeks.

### For the sotorasib combination studies

with either copanlisib (6 mg/kg) or sapanisertib (1mg/kg) in the TC314AR PDX model, treatment arms included single-agent administration of each agent to assess the contributions of each agent. Control mice were treated with the solvent. Sotorasib, copanlisib or sapanisertib, or solvent were administered QD orally for 21 days. Tumor growth was monitored 2-3 times/week.

The following formula approximates the volume of the ellipsoid corresponding to the tumor: tumor volume (mm^3^) = [tumor length X (tumor width)^2^]/2. Tumor growth curves show median or average tumor volumes, and the standard error (SE) or standard deviation was added to demonstrate variability. Changes in tumor volume were assessed by comparing the volume at a defined time point with the baseline volume, thus adjusting for baseline tumor volume. For an individual mouse, the tumor response according to the percent change in tumor volume was calculated from baseline to the time of assessment as follows: % tumor volume change *ΔVt* = (*Vt -V0/ V0*) X *100*, where *Vt* is the tumor volume at time *t*, and *V0* is the tumor volume at baseline. We used the following criteria to determine treatment responses: 1) Tumor Regression (or partial response): tumor regression ≥ -30% based on tumor volume changes calculated by AUC_0-21day_ or at day 21 after treatment start when compared with baseline (beginning of treatment at day 0); 2) Tumor Growth Inhibition (or stable disease): Tumor growth was significantly suppressed when compared with control (p < 0.05), but no tumor regression was observed, or tumor regression was less than -30% based on tumor volume changes calculated by AUC_0-21day_ or at day 21 after treatment start when compared with baseline; 3) Resistance: Tumor volume changes calculated by AUC_0-21day_ was not significantly different from control group (p>0.05).

### Development of sotorasib-acquired resistant PDX models

To develop NSCLC PDXs with acquired resistance to sotorasib (AMG510), *KRAS*^*G12C*^ mutation harboring TC303 and TC314 PDXs, the two most sotorasib-sensitive PDXs, were implanted into NSG mice. when the tumor volume of these PDXs reached approximately 200 mm^3^, mice were treated with AMG510 (100mg/Kg) daily for at least 3 weeks, paused if the tumor size shrank below 200 mm^3^. We monitored mice with regressed tumors for tumor regrowth. When those tumors regrew to 200 mm^3^ in size, we retreated again, until mice were euthanized at a tumor size of around 500-100 mm^3^. The tumors were passaged to new NSG mice for another round of AMG510 treatment. Third-generation (G3) PDXs TC303AR and TC314AR were generated, in which each generation was treated with three or more cycles. Susceptibility to AMG510 (sotorasib) was reduced in each passage. We performed whole-exome sequencing for at least 2-3 tumors from each PDX obtained in passage 3 (G3) that were under constant AMG510 treatment and did not regress during treatment. All resistant tumors from the third generation from both PDXs had the same *KRAS*^*G12C*^ mutation as the parental PDXs (TC303 and TC314), with no other *KRAS* mutations.

### PDX-derived organoid (PDXO) culture, maintenance, and development of sotorasib-acquired resistant PDXO models

PDX tissues were harvested and processed into 2-3-mm-diameter pieces and washed with ice-cold PBS. Tumor pieces were dissociated into single cells in Advanced DMEMF12 (GIBCO) with Liberase TM (Sigma) for 1 hour, followed by a 10-minute incubation with TrypLE Express (Invitrogen) at 37^0^C with gentle shaking. The establishment of organoids in culture was done according to the protocol described in Shi R et al (46) with little modification. Briefly, single cells from freshly harvested PDX tissues were counted and resuspended in 100% growth factor–reduced Matrigel (VWR), plated in 24-well tissue culture plates as Matrigel domes, and maintained in 37^0^C 5% CO2 with media overlaying the Matrigel dome. Growth factors and supplements used in the media for the organoid culture are followed according to the reference article. Organoid growth was monitored weekly for the detection of initiated organoids, and organoids were kept in the same passage for no longer than four weeks. To make the sotorasib-resistant PDXOs, PDX TC303 and PDX TC314 tissues were harvested, and a single-cell suspension and establishment of organoids (PDXO303 & PDXO314) were prepared according to the protocol mentioned above. These organoids were cultured in a complete medium with all supplements and growth factors and incremental doses of sotorasib for 1-4 weeks before passaging. Repeated these passaging steps with sotorasib for several cycles until the organoids grew smoothly without cell death with 1 µM sotorasib-containing medium and developed the sotorasib-resistant PDXO303AR and PDXO314AR isogeneic organoids. The resistant organoids were then subjected to whole-exome sequencing to verify *KRAS*^*G12C*^ mutation status.

To check the purity of organoids, we prepared the single cells from fresh organoids and stained them with a mouse cell detecting antibody (H-2Kb/H-2Db; BioLegend, CA, USA) and a human epithelial marker (EpCAM; BioLegend, CA, USA) and ran flow cytometry using Attune NxT cytometer (Invitrogen, CA, USA). Data were analyzed by Flow Jo software. Organoid cultures were tested routinely for Mycoplasma. The established organoid samples were subjected to whole-exome sequencing (method mentioned above) for detecting *KRAS*^*G12C*^ mutation.

### Generation of 4E-BP1 and PI3K/p110a knockout clones by CRISPR-Cas9 technology

CRISPR-Cas9 technology was used to knock out PI3K and 4E-BP1 genes so that the corresponding functional proteins are no longer produced. These experiments involved transfecting H23AR and H358AR cells with a target-specific guide RNA (gRNA) and a Cas9 nuclease. All the reagents, which included Gene Knockout Kit v2, sgRNA, and SpCas9 Nuclease, were purchased from Synthego (Synthego, CA, USA). The experiment was performed according to the manufacturer’s protocol. Briefly, for one knockout test, 3 µl sgRNA (30 µM), 0.5 µl Cas9 (20 µM), and 3.5 µl Buffer R were mixed and kept at Room Temperature for 10 min, and then 1 x 10^5^ cells in 3 µl Buffer R were added to the above mixture, followed by electroporation (Invitrogen Neon electroporation system: 10-µl tip type) (Pulse voltage, 1100 V; Pulse width, 30 ms; Pulse number, 1). After electroporation, the knockout cells were cultured in a 24-well plate at 37 ^0^C for 3 days. Total genomic DNA was extracted, PCR for specific genes was conducted, a certain DNA fragment was sequenced with the Sanger technique, and sequencing results were analyzed using Inference of CRISPR Edits (ICE) (a free online tool on the Synthego website) that provides an easy quantitative assessment of indels generated by CRISPR in pools and clones. The ICE report outputs both the total indel percentage and the Knockout Score (the percentage of outcomes that lead to a putative knockout). Finally, the knockout efficiency was carried out using a Western blot for the specific knockout gene protein products.

### Cell viability assay in organoids (PDXOs)

Organoids were dissociated into single cells, counted, and prepared a cell suspension with media containing Matrigel and plated cells in 96-well u-shaped plates (5000-10000 cells per well) in triplicate for 24-72 hours before drug treatment. Organoids were treated with a range of drug concentrations (0.01– 10 mmol/L) for 7 days, and cell viability was determined by Cell Titer Glo 3D viability assay according to the manufacturer’s protocol (Promega, WI, USA). Drug–response curves were graphed, and IC50 values were calculated using GraphPad Prism 10.0. CompuSyn software (46) was used to calculate the combination indices for combination drug studies.

### Drug sensitivity assay

Sotorasib-resistant isogenic cells were seeded at a density of 3X10^3^ cells/well in a 96-well microplate and treated with sotorasib or adagrasib at concentrations ranging from 0.001 to 10 µM in DMSO. Cells were incubated in 37°C incubators, 5% CO_2_, for three days. Cytotoxicity assays were performed using colorimetric XTT (Sigma-Aldrich, USA) and SRB (Sulforhodamine B) Assay Kit (Abcam, USA) reagents according to the manufacturer’s protocol. Optical density (OD) was measured using a microplate reader (FLUOstar Omega, BMG Labtech USA) at 570 nm. Cell viability (%) = [OD (Drug) - OD (Blank)] / [OD (Control) - OD (Blank)] × 100.

### Colony formation assay

NCI-H23, NCI-HH23AR, NCI-H358, and NCI-H358AR cells were seeded into 6-well plates at a density of 300 cells per well, and treated with copanlisib alone, MK2206 alone, copanlisib + MK2206 combination, everolimus alone, and copanlisib + everolimus combination at the concentrations listed in the experiment for 72 h. The media was replaced every 2-3 days with medium containing drugs. After twelve days, colonies were fixed with 4% PFA, stained with crystal violet solution, and photographed.

### Western blot analysis

Cells were seeded at a density of 3X 10^6^ in 90 mm Prime Surface plates (Corning), incubated overnight in RPMI1640 complete medium, treated with appropriate inhibitors, and harvested. Western blotting was performed according to the manufacturer’s protocol. Total protein was harvested using Ripa lysis buffer (Merck, Burlington, MA), and its concentrations were evaluated with BCA™ protein assay kit (Pierce, Rockford, IL, USA). Equal amounts of proteins were separated by 8-15% SDS-PAGE gel, electro-transferred onto a Hybond ECL transfer membrane (Amersham Pharmacia, Piscataway, NJ), and blocked with 2-5% non-fat skim milk. Then, membranes were probed with specific primary antibodies at 1:1000 dilution overnight at 4 °C, washed with PBS, and incubated with corresponding secondary antibodies at 1:2000 dilution at room temperature for 1 h. The specific protein bands were visualized with an ECL advanced western blot analysis detection kit (GE Health Care Biosciences, NJ, USA). Antibodies used in this study were purchased from Cell Signaling (Beverly, MA): Anti-PI3K(p110a) (#4249S), PI3K (p85), AKT (CS#4691), p-AKT (Ser473) (CS#9271), p-AKT (Thr308) (CS #4056), mTOR, p-mTOR (Ser2448) (CS#2971), PDK1 (CS#13037), p-PDK1(Tyr373/376) (bs-3017R), p-PDK1(Ser241) (CS#3061), MAPK (CS#4695), p-MAPK (Thr202/Tyr204) (CS#4370), PTEN (CS#9559), p-PTEN (CS#9549), p-S6 (Ser240), p-S6 (Ser235), 4E-BP1 (#9644S), p4E-BP1 (Ser65) (#9456S), p4E-BP1 (T37/46), p-PRAS40 (Thr246), p-GSK-3b (Ser9), p70S6 (T389). Monoclonal anti-β-actin (Sigma#A5449) was purchased from Sigma Aldrich (St Louis, MO). GAPDH mouse monoclonal antibody (sc-365062) was from Santa Cruz Biotechnology (CA, USA).

### Whole Exome Sequence

H23 & H23AR and H358 & H358AR isogeneic cells were seeded in triplicate at a cell density of 2 X 10^6^/plate. DNA was isolated and purified using a Qiagen kit (Germantown, MD). DNA was also isolated from acquired resistant PDXO303AR, PDXO314AR organoids, as well as TC303AR & TC314AR PDXs with their sensitive counterparts for whole exome sequencing. The quality of DNA was evaluated, and the whole exome was sequenced at the sequencing core lab (MDACC, Houston, TX) using a next-generation sequencer (NextSeq500, Illumina, USA). Sequencing data were analyzed by the Department of Bioinformatics and Computational Biology, MDACC.

### Reverse Phase Protein Array (RPPA)

NSCLC PDXs treated with sotorasib, and the residual PDX samples were harvested and snap-frozen for RPPA analysis. Similarly, H358AR CDX tumors were treated with sotorasib, copanlisib, and their combination. After the experiments, the residual tumor tissues were harvested, snap-frozen, and stored at -80^0^C. RPPA analysis was performed using 400 antibodies at the RPPA core lab at UT MD Anderson Cancer Center. Bioinformatics analysis was performed by the Department of Bioinformatics and Computational Biology at MD Anderson Cancer Center.

### Mass Spectrometry

Sotorasib-acquired resistant PDXs (TC303AR and TC314AR) and their parental counterpart PDXs (TC303 and TC314) were submitted to the core lab in Baylor College of Medicine (BCM) (Houston, TX) for mass spec analysis. Three biological replicates were used in each PDX. The samples were denatured and lysed by three cycles of LN2 snap freeze and thaw at 95 °C. For global profiling, 10 mg of lysate was trypsinized to obtain 10 mg of digested peptides. After fractionation using a small-scale basic pH reverse phase (sBPRP) step elution protocol with increasing acetonitrile concentrations, fractions were combined into 5 pools that were resolved and sequenced online with Fusion Lumos and timsTOF fleX mass spectrometer. For phospho-proteome profiling, a 100 µg protein lysate was digested with trypsin and dried under vacuum. Global and phospho-proteomic analyses covered over 8000 gene protein products (GPS), and over 4000 GPS, respectively, which included the kinome profile. After label-free nanoscale liquid chromatography coupled to tandem mass spectrometry (nanoLC-MS/MS) analysis using a Thermo Fusion Mass spectrometer, the data were processed and quantified against NCBI RefSeq protein databases in Proteome Discover 2.5 interface with a Mascot search engine (Saltzman, Ruprecht). The Skyline program was used to obtain precise quantification. To decipher phospho-proteome signal pathway analysis, we utilized protein external data contributions for phosphorylation-related data mining sets, including PhosphoSitePlus (http://www.phosphosite.org/), Phospho.ELM, PhosphoPep, and the Phosphorylation Site Database (PHOSIDA).

## Statistics and Reproducibility

### In-vivo xenograft, CDX, and PDX data analysis

All in-vivo experiments were designed and planned with biostatistician Dr. Jing Wang (Co-author) from the bioinformatics department at MD Anderson Cancer Center. For therapeutic studies in subcutaneous xenografts, CDX or PDX tumors were developed in immune-deficient NSG mice, and generalized linear regression models were used to study the tumor growth over time. Statistical analyses were performed with GraphPad Prism 7 software. Tumor volume change per time point was calculated as a relative level of tumor volume change from baseline. Two-way ANOVA with the interaction of treatment group and time point was performed to compare the difference in tumor volume changes from baseline between each pair of the treatment groups at each time point. Means ± standard errors of the mean are shown in all graphs. The nonparametric Mann-Whitney U test was applied to compare cell numbers in different treatment groups. Differences of *P* < 0.05, *P* < 0.01, and *P* < 0.001 were considered statistically significant. All in-vivo tumor growth and treatment experiments were repeated N =3 times with N = 5-7 mice/group each time.

### Combination Index analysis to determine treatment synergy

We evaluated the potential synergistic effect of the drug combinations sotorasib + copanlisib or sotorasib + sapanisertib under the Highest Single-agent (HSA) framework (47), where the synergistic effect of a drug combination is declared if the combination effect is greater than that of the more effective individual component. The combination index (CI) under the HSA and the corresponding standard error were approximated by the Delta method (48). We considered the three most well-established reference models: Highest Single Agent (HAS), response additivity (RA), and Bliss Independent (BI), in a unified statistical quantification method (48). The in-vivo experiment was repeated N =3 times with at least N = 5 mice/group at each time.

#### Reverse Phase Protein Array (RPPA) data analysis

Slides were scanned using a CanoScan 9000F, and spot intensities were quantified using ArrayPro Analyzer 6.3 (Media Cybernetics, Washington, DC). SuperCurve, a software developed in-house, was used to estimate relative protein levels (49). After SuperCurve fitting, protein measurements were normalized for loading using median centering across antibodies. One-way analysis of variance (ANOVA) was used to assess the differences in protein expression between control and treatment groups on a protein-by-protein basis. First, for one feature(protein) at a time, we carried out an overall F test to detect any significant difference among the means of all the groups. Next, for the featured (proteins) identified in this process, we then compared between desired groups to identify the sources of difference. The R library “multcomp” was used for this purpose. Note that the FC (fold change) values were calculated as the estimated ratio between the 2 groups in comparison, with the following conventional modification: For the rations > 1 (up-regulation), FCs were noted as the same as the ratio. For the ratios ≤ 1 (down-regulation), FC were noted as the negative inverse of the ratio. Furthermore, to account for multiple testing, we estimated the false discovery rates (FDR) of the overall test of the model using the Benjamini-Hochberg method. An FDR-adjusted p-value less than 0.05 was considered statistically significant unless otherwise mentioned. The criteria of significant protein selection were: 1. Significant in overall F-test (FDR-adjusted p-value<0.05); 2. Significant in pairwise comparison (FDR-adjusted p-value<0.05). N=3-5 tumor samples per group were used for RPPA analysis.

#### Mass Spectrometry data analysis

Statistical analysis was performed using R software (R version 4.0.1). The log2 transformation was applied to the iFOT Half Min proteomic data. The Student’s t-test was used to compare expression values between the groups. P values obtained from multiple tests were adjusted using FDR. Statistical significance was defined as FDR < 0.05. The enriched pathways and hallmarks were identified by pre-ranked GSEA using the gene list ranked by log-transformed P □ values with signs set to positive/negative for a fold change of >1 or <1, respectively. N=3 samples/group were analyzed for Mass spectrometry.

### Whole Exome Sequence data analysis

The quality of raw FASTQ reads was assessed using FastQC and then mapped to the human reference genome GRCh38, using BWA (50,51). The reference genome refers to the b38 version with decoy sequences for human GRCh38 provided in the Genome Analysis Toolkit (GATK) resource bundle (52). The mutations were called following the GATK best practice pipeline. The candidate mutations were filtered for high-confidence somatic mutations and annotated for functional changes using ANNOVAR (53). At least N=3 samples/group were analyzed for WES.

### Cell Survival Assay data analysis

The percentage of viable cells was determined by the ratio of absorbance of treatment and control groups: ODT/ODC x 100%. Univariate analysis was performed to evaluate the distribution of data for each treatment group. To determine whether SRB% % was different between treatment groups, two methods were used: 1) ANOVA was performed to compare the variance between treatment groups for all samples within each cell line, and 2) Tukey’s multiple comparisons test was performed for pairwise differences between treatment groups. P<0.05 was considered statistically significant; all tests were two-sided. Analyses were performed using SAS 9.3 (SAS Institute Inc., Cary, NC). Values represent the mean of three independent experiments. Each assay was performed N ≥ 3 times with at least 3 replicates each time.

## Data Availability Statement

All data generated or analyzed during this study are included in this published article. The source data underlying most graphs and charts used in this manuscript will be provided as Excel files in Source Data 1. All uncropped and unedited Western blot images used in this manuscript will be provided as supplementary information in Supplementary Figures. Requests for any source data or materials that are not provided should be made to the corresponding author.

## Acknowledgments

We acknowledge the MD Anderson animal facility for providing a dedicated housing room for the mouse study; the MD Anderson RPPA core lab for RPPA experiments; the MD Anderson NSCLC PDXNet core; the MD Anderson Department of Scientific Publication, BCM Mass Spectrometry core, and Dr. Yonathan Lissanu for his suggestions in lab meetings.

This work was supported in part by the National Institutes of Health/National Cancer Institute through The University of Texas MD Anderson Cancer Center’s Cancer Center Support Grant (CCSG) CA-016672 - Lung Program and Shared Core Facilities (J.A. Roth); Specialized Program of Research Excellence (SPORE) Grant CA-070907 (J. A. Roth); PDX development and trial grant U54CA-224065 (J. A. Roth); U54 program grant on Acquired Resistance, ARTNet U54CA224081 (J.A. Roth); Lung Cancer Moon Shot Program (J.A. Roth), The University of Texas MD Anderson Cancer Center, sponsored research agreement from Genprex, Inc.

## Figure legends

**Figure 1-figure supplement 1:**
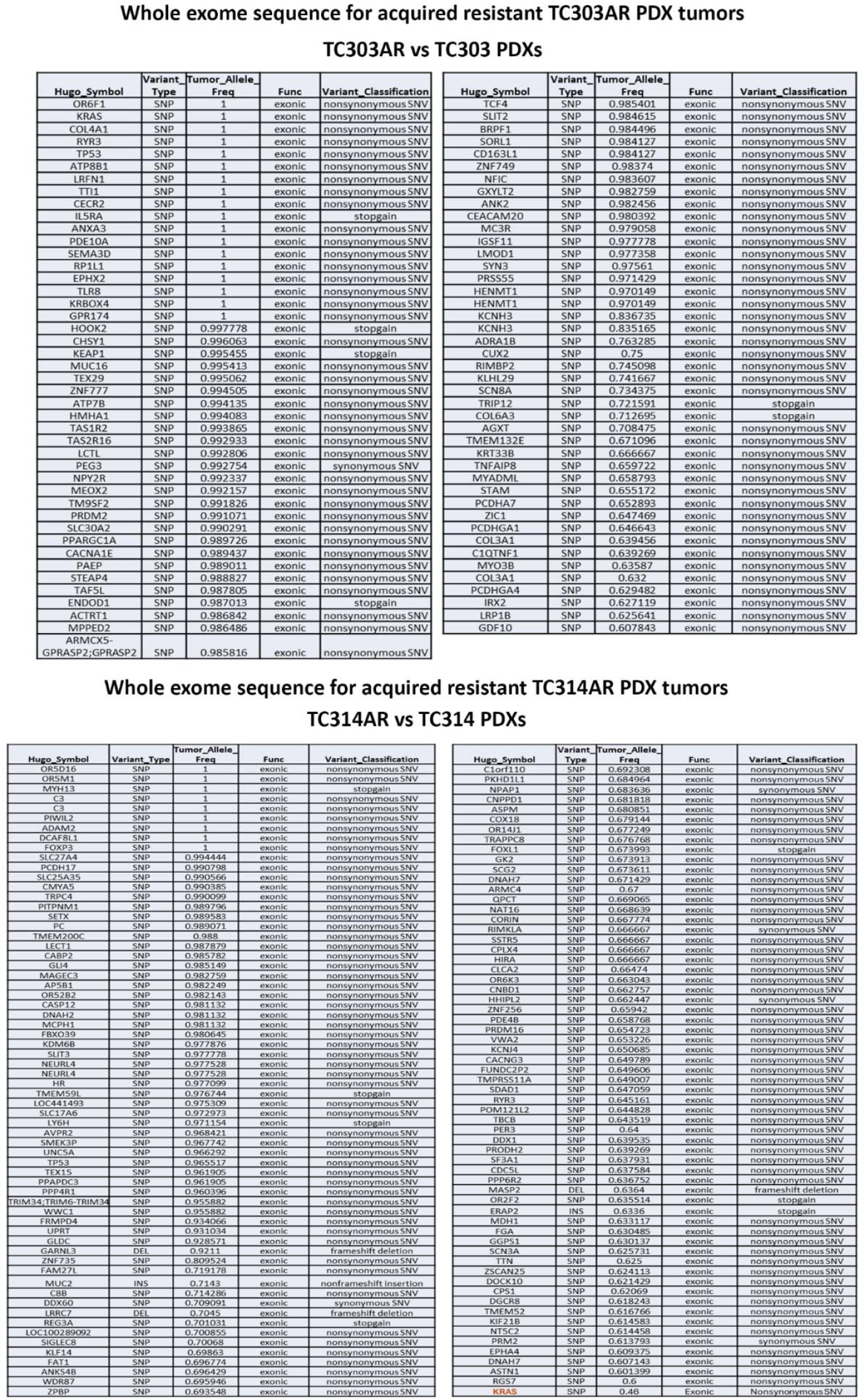
Whole exome sequence (WES) on acquired resistant PDXs. Sotorasib-resistant TC303AR & TC314AR PDXs were subjected to WES to identify additional mutations in KRAS as compared with their parental TC303 & TC314 PDXs. Three biological replicates for each sample were used.

**Figure 2-figure supplement 1:**
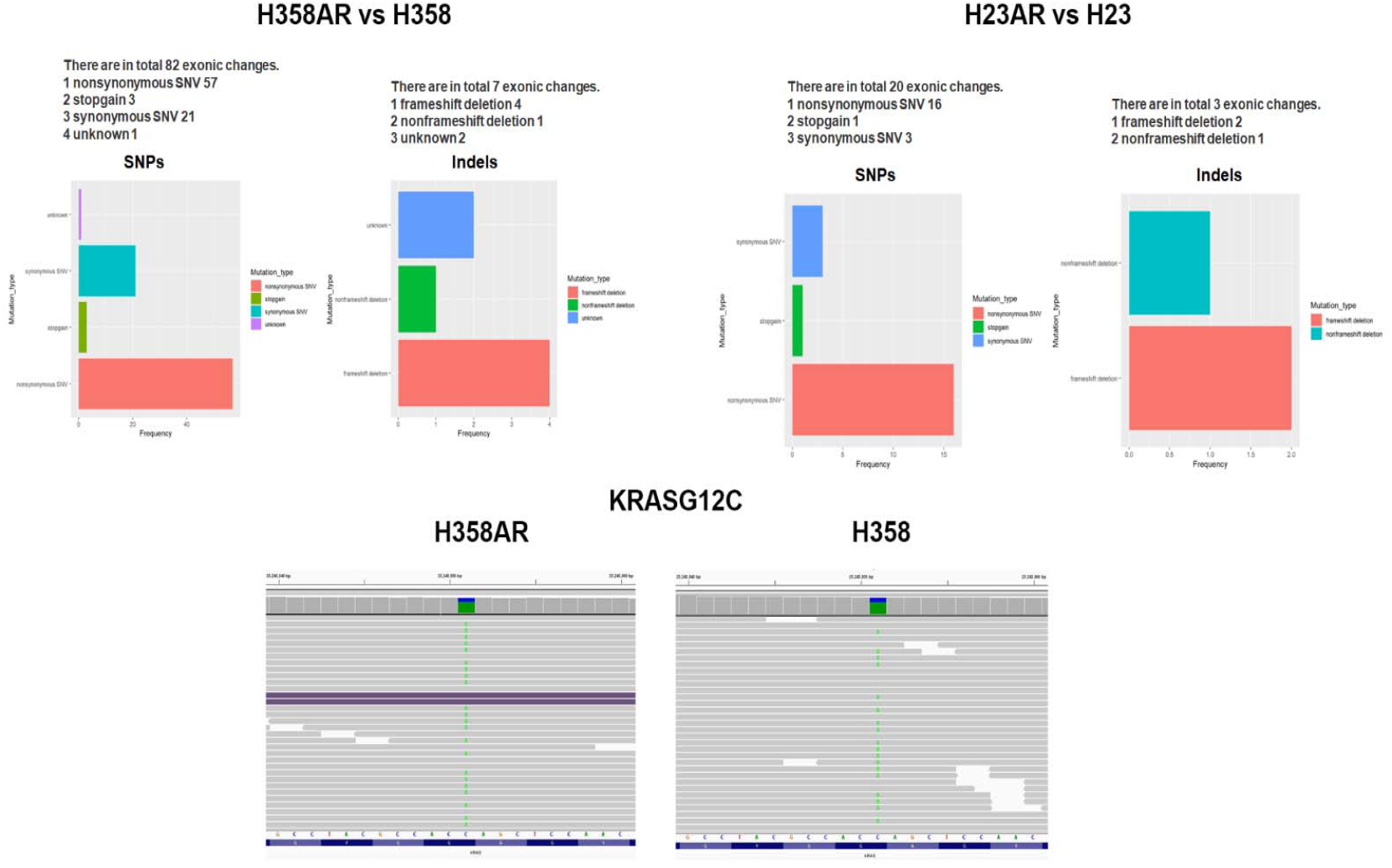
Whole exome sequence (WES) on acquired resistant cells. Two sotorasib-resistant isogeneic cell lines were subjected to WES to identify additional mutations in *KRAS*. Three biological replicates for each sample were used. Top left: number of SNPs and Indels found in H358AR as compared with the parental H358 cells. Top right: SNPs and Indels found in H23AR cells as compared with H23 cells. Bottom: *KRAS*^*G12C*^ mutation is shown to be present in both H358 and H358AR cells.

**Figure 3-figure supplement 1:**
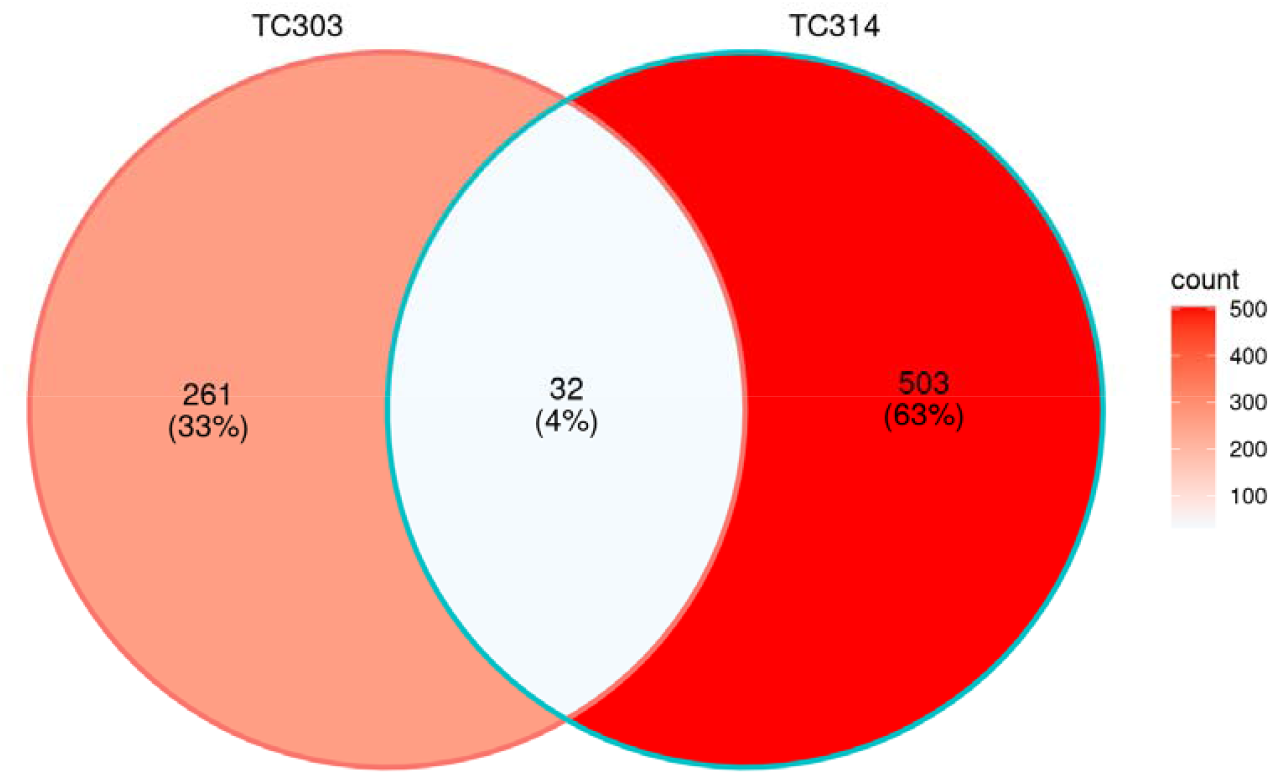
Venn Diagram. Mass spectrometry was performed for two sotorasib-resistant isogenic PDXs (TC303AR and TC314AR) samples. Altered proteins from each pair of isogenic PDXs were used for the Venn Diagram. The percentage of commonly altered proteins is shown in the middle (white area).

**Figure 3-figure supplement 2:**
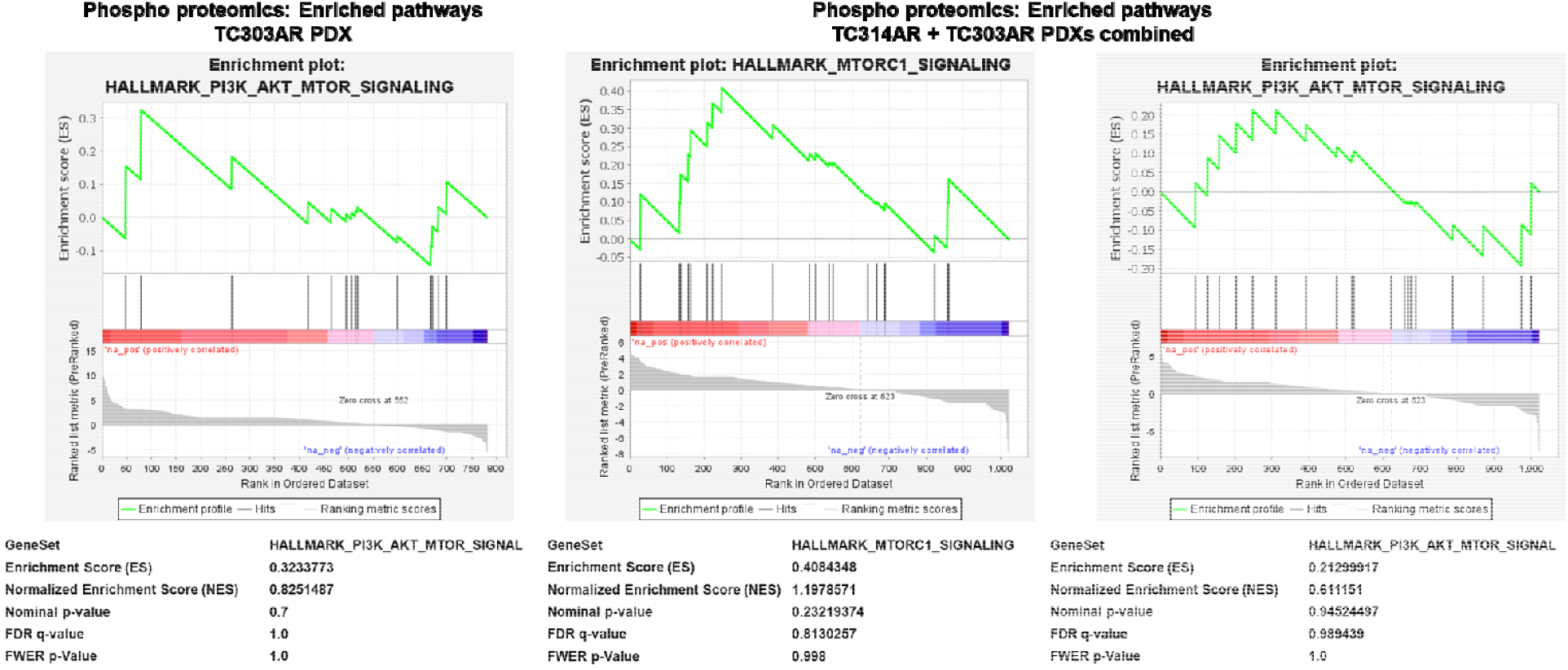
Mass spectrometry analysis for phosphoproteomics in resistant PDXs. Mass spectrometry was performed for two sotorasib-resistant isogenic PDXs (TC303AR and TC314AR) samples. Phosphoproteomics data were analyzed for enrichment pathway analysis. Enrichment graph for PI3K/AKT/mTOR signaling in TC303AR PDX (Left) and Enrichment graphs for mTORC1 and PI3K/AKT/mTOR signaling in TC314AR+TC303AR PDXs combined data analysis (middle and right).

**Figure 3-figure supplement 3:**
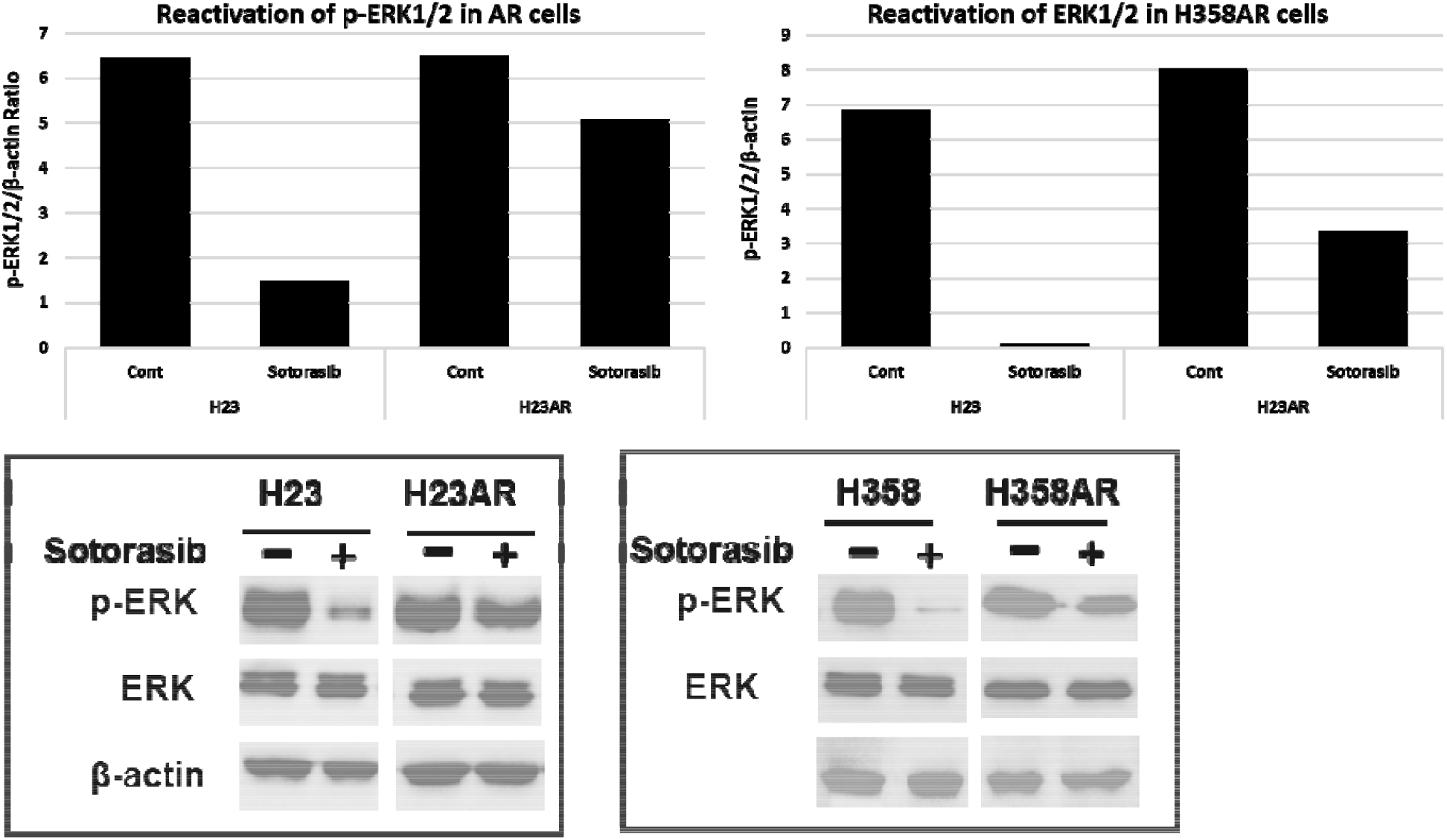
KRAS Reactivation in Sotorasib-Resistant Cells via ERK Phosphorylation. H23, H23AR, and H358 & H358AR cells were compared for their p-ERK1/2 expression. Both H23AR and H358AR resistant cells were cultured and maintained under sotorasib pressure. A). Western blots for total ERK and p-ERK in these cells are shown. B) Quantitation of p-ERK expression in AR cell lines and compared with their parental counterpart.

**Figure 4-figure supplement 1:**
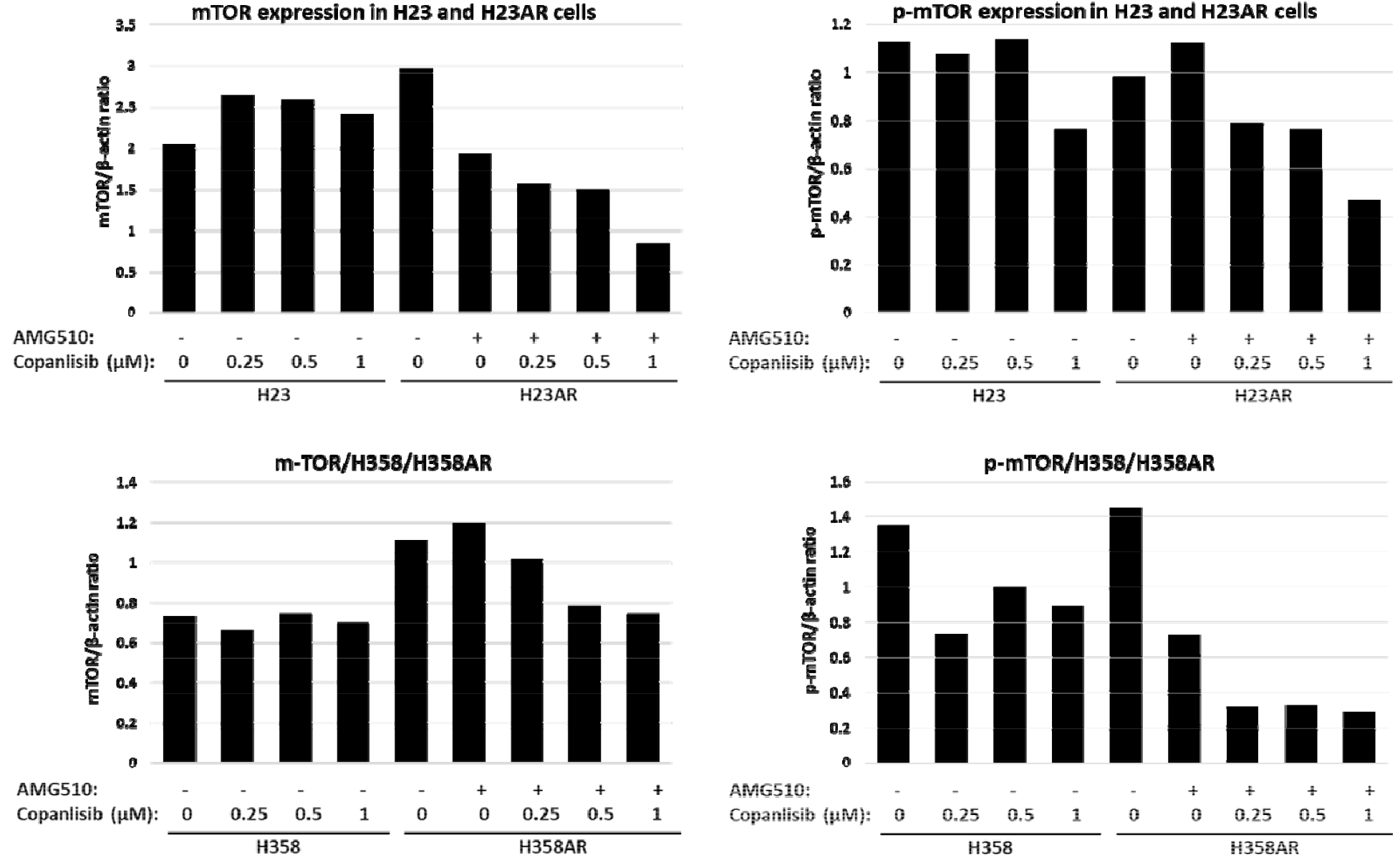
Effect of copanlisib on mTOR and p-mTOR expression. H23, H23AR, and H358 & H358AR cells were cultured. Both sotorasib-resistant H23AR and H358AR cells were maintained under sotorasib pressure during culture; all cells were treated with different concentrations of copanlisib for 24 hr. Western blots for mTOR and p-mTOR were performed and shown in Figure 4G. The quantitative analysis for these proteins was done and compared between resistant vs sensitive cells; H23 vs H23AR and H358 vs H358AR cells.

**Figure 4-figure supplement 2:**
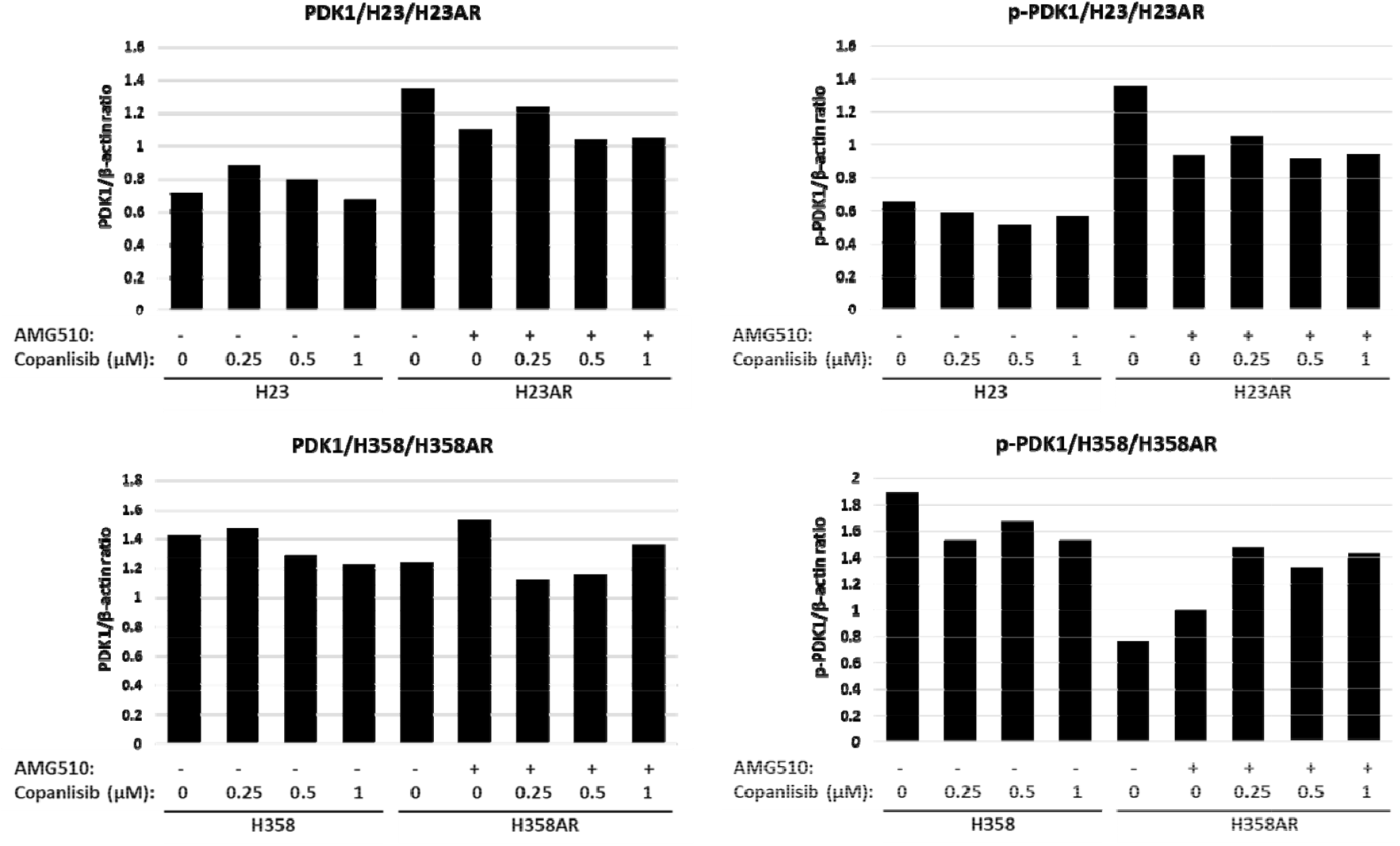
Effect of copanlisib on PDK1 and p-PDK1 expression. H23, H23AR, and H358 & H358AR cells were cultured. Both sotorasib-resistant H23AR and H358AR cells were maintained under sotorasib pressure during culture; all cells were treated with different concentrations of copanlisib for 24 hr. Western blots for PDK1 and p-PDK1 were performed and shown in Figure 4G. The quantitative analysis for these proteins was done and compared between resistant vs sensitive cells; H23 vs H23AR and H358 vs H358AR cells.

**Figure 7-figure Supplement 1A.**
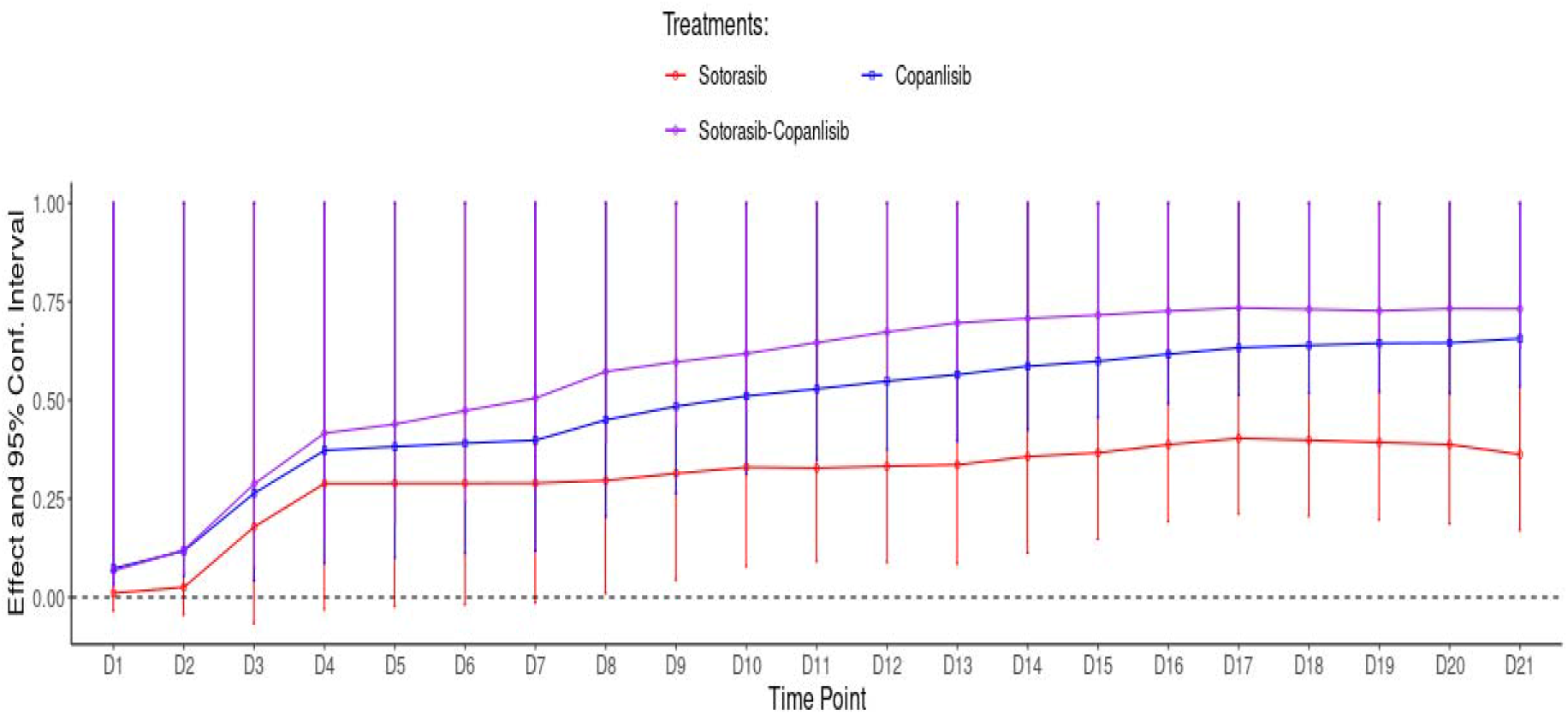
Treatment effect on TC314AR PDX by sotorasib + Copanlisib combination treatment. The resistant TC314AR PDX tumors were treated with the combination and single agents. The antitumor effect data were analyzed by CombPDX analysis software to determine the treatment effect. The treatment effect is measured as the percentage of tumor growth inhibition with a 95% confidence interval.

**Figure 7-figure Supplement 1B.**
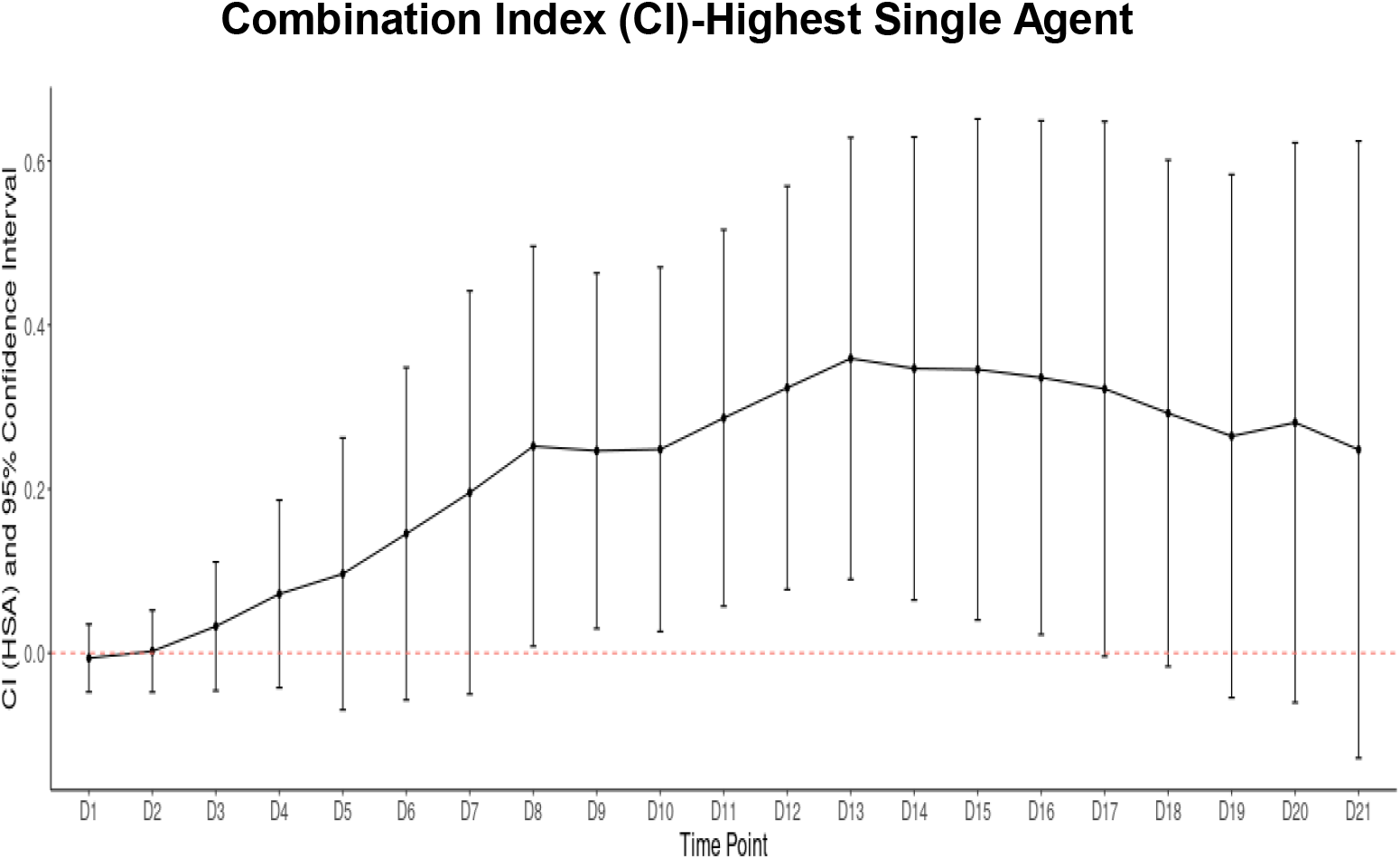
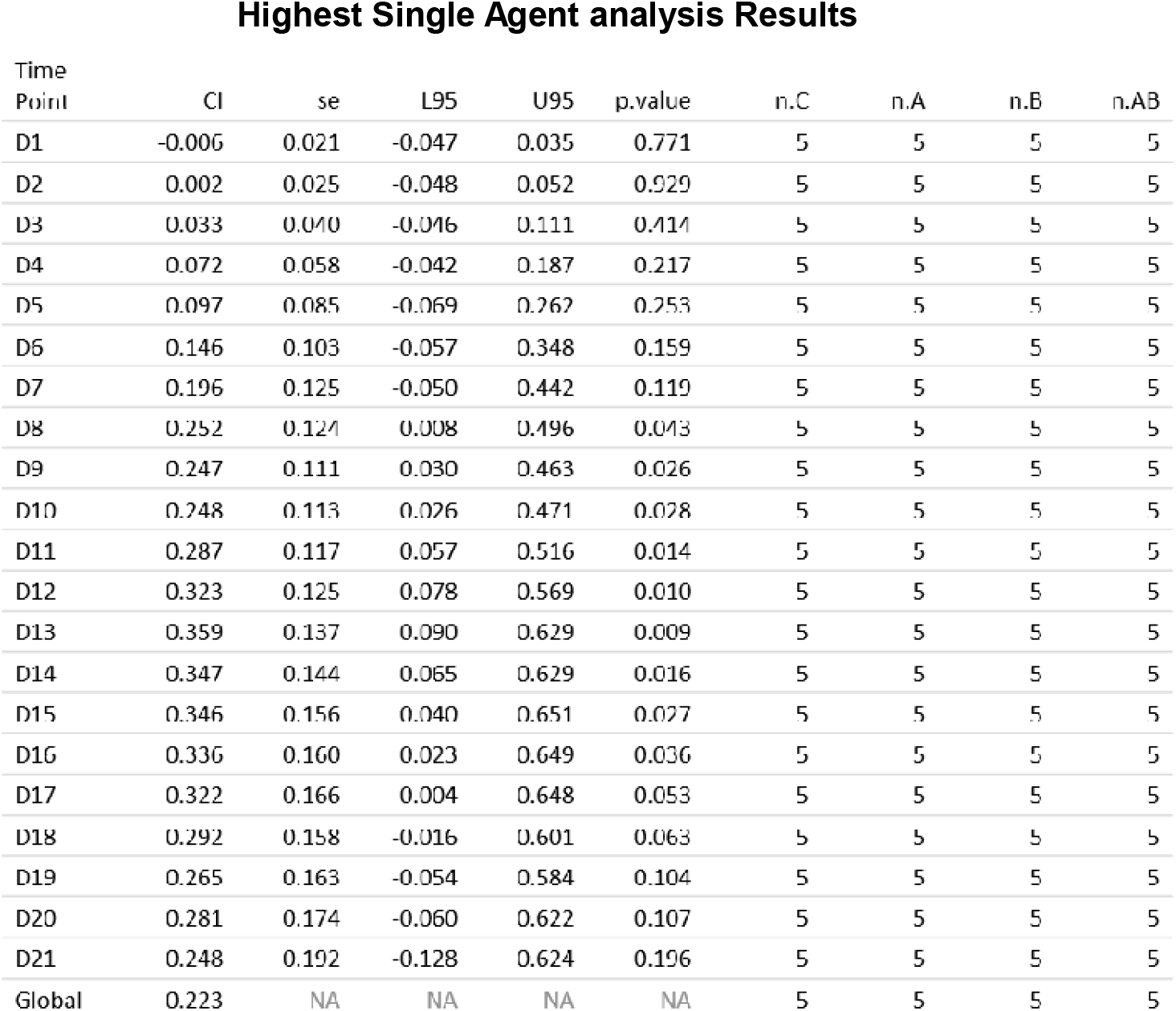
The analysis of the synergistic effect of sotorasib + copanlisib combination treatment on TC314AR PDX tumors. The resistant TC314AR PDX tumors were treated with the combination and single agents. The synergistic antitumor effect was calculated by analyzing the combination index (CI) using CombPDX analysis software to determine the synergy between these two drugs. The highest single agent method with a 95% confidence interval was used to determine the combination index (CI). CI values <1 are synergistic.

**Figure 7-figure Supplement 2A.**
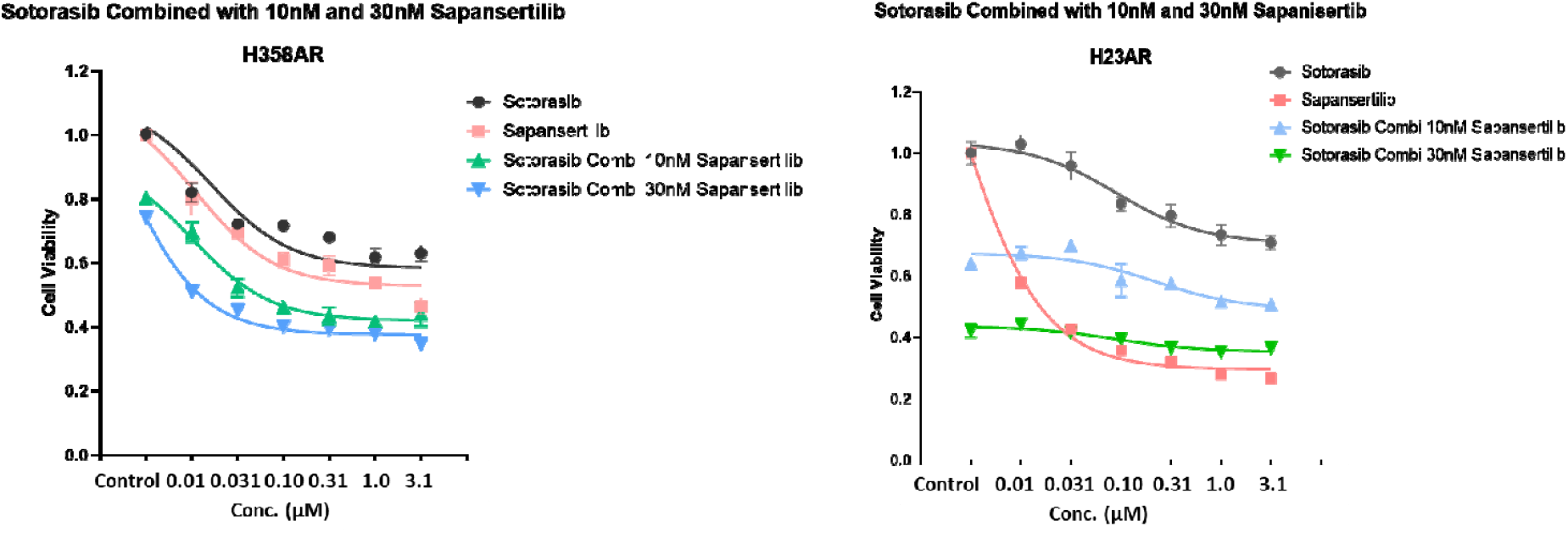
Cell viability assay for sotorasib + sapanisertib combination on H23AR and H358AR cells. The experiment was done three times. Data is shown as mean percentage ±SD, n=3

**Figure 7-figure Supplement 2B.**
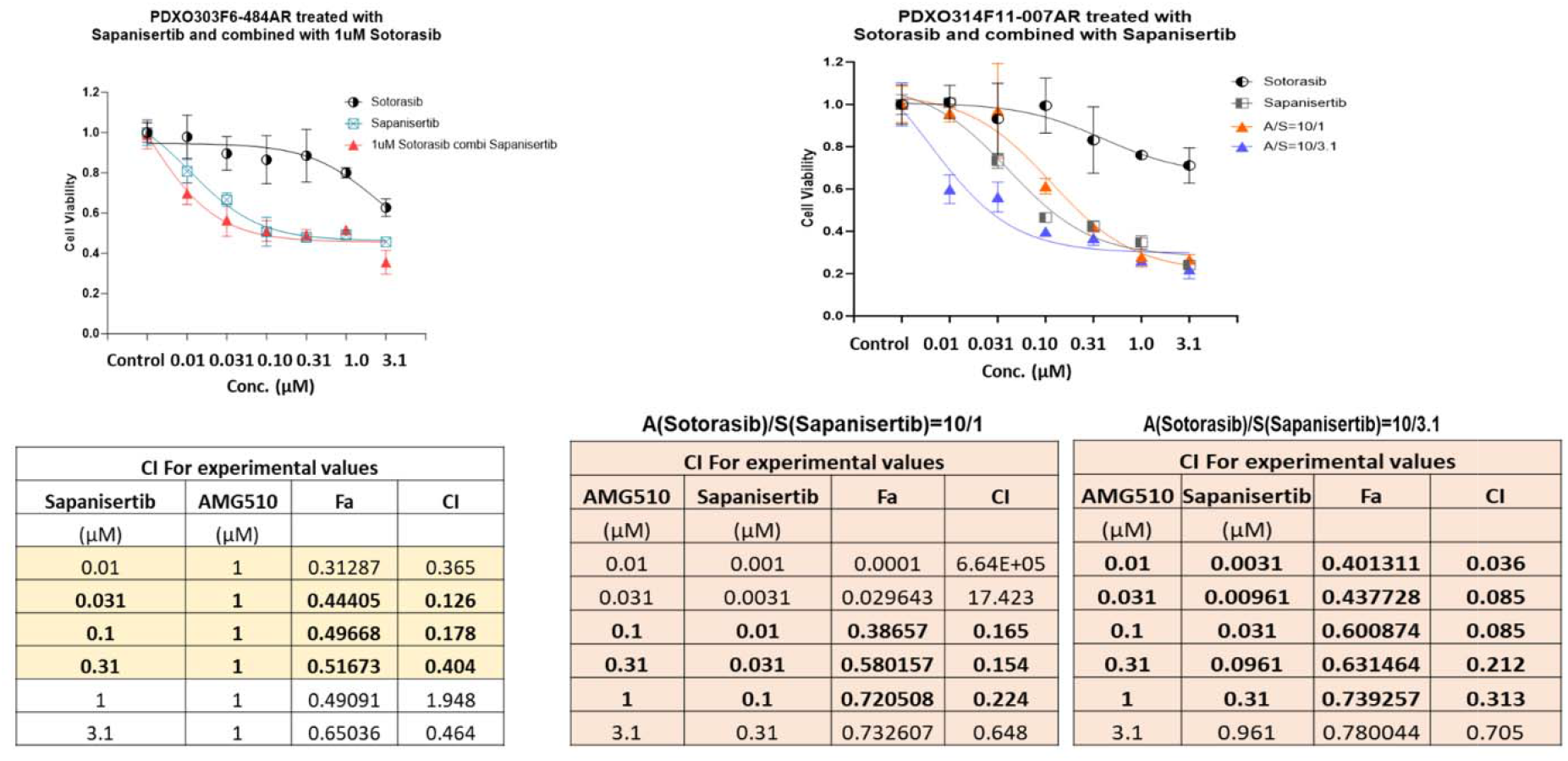
Cell viability assay for sotorasib + sapanisertib combination on PDXO303AR and PDXO314AR organoids. The organoids were treated with sotorasib in combination with sapanisertib. The combination index (CI) for cell viability was calculated using the CalcuSyn software using the Chou-Talalay method. If the CI value is <0.7, the combination is synergistic; if the CI value is between 0.3-0.1, the combination is strongly synergistic, and if the CI value is <0.1, the combination is very strongly synergistic. The experiment was done three times. Data is shown as mean percentage ±SD, n=3

